# Saxiphilin is a broad-spectrum toxin sponge for C13-modified saxitoxins

**DOI:** 10.64898/2026.03.10.710854

**Authors:** Sandra Zakrzewska, Zhou Chen, Elizabeth R. Park, Roshni G. Bhaskar, T. Aaron Bedell, J. Du Bois, Daniel L. Minor

## Abstract

Saxitoxin (STX) and its congeners (paralytic shellfish toxins, PSTs) are among the most potent small-molecule toxins. PSTs are produced by harmful algal blooms and derive toxicity by disrupting voltage-gated sodium channel (Na_V_) bioelectrical signaling. Understanding how PST structural variation affects target binding is crucial to develop means to counteract PSTs and exploit these natural products as drug development leads. Frog and toad saxiphilins (Sxphs) are soluble, high-affinity STX toxin sponge proteins that offer a powerful platform to define PST-protein interactions. Here, we show that American bullfrog (*Rana catesbeiana*) *Rc*Sxph and High Himalaya frog (*Nanorana parkeri*) *Np*Sxph bind a broad set of C13-modified STX congeners. High-resolution X-ray crystal structures of toxin complexes with *Rc*Sxph, *Rc*Sxph mutants, and *Np*Sxph unveil two C13-aryl congener binding modes, termed ‘compact’ and ‘open’, that depend on the *Rc*Sxph Tyr558 local environment. These results highlight a remarkable adaptability of Sxphs for accommodating chemically diverse STX analogs and reveal unexpected toxin conformational plasticity. These findings have implications for understanding PST interactions with biological targets and informing design of STX-based probes and Na_V_ modulators.

## Introduction

Saxitoxin (STX) and its congeners represent a major class of paralytic neurotoxins that pose severe risks to human health through paralytic shellfish poisoning (PSP) resulting from ingestion of contaminated seafood^1–5^. This family of lethal neurotoxins, called paralytic shellfish toxins (PSTs), is produced by marine dinoflagellates and freshwater cyanobacteria that cause harmful algal blooms (HABs)^1,3,4^ and are potent inhibitors of bioelectrical signal transmission through voltage-gated sodium channels (Na_V_s) in nerve and muscle^1,3^. PSTs comprise >50 STX congeners, commonly encompassing modifications of the STX scaffold at three sites (Fig. 1A): R1, the carbamoyl site; R2, the six membered guanidinium ring N1 position; and R3, the C11 carbon^1,2,6^. Due to the global impact of PSP and HABs^3,5^, classification of STX as a chemical weapon^7^, and lack of an effective PSP antidotes^4^, there is an ongoing effort to define how STX and its congeners engage target proteins^8–12^ and to develop STX-neutralizing agents^13–17^. Beyond their toxic effects, STX and its derivatives are exploited as powerful research tools for probing Na_V_ function and physiology^18–21^ and, despite their reputation as toxins, hold pharmaceutical potential^22–25^. Hence, understanding the molecular recognition rules governing various STX forms^10,12,19,21^ remains an important goal for developing anti-PSP strategies and creating new precision small molecules to control Na_V_ signaling.

**Figure 1.**
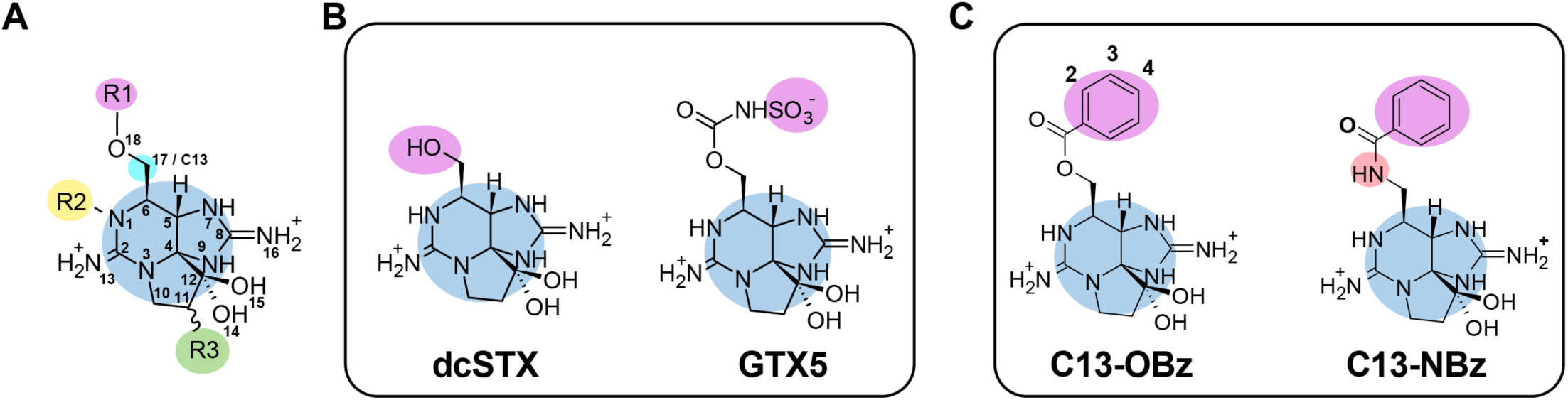
C13-modified STX congener examples. **A**, STX core (blue) showing modification sites, R1 (magenta), R2 (yellow), and R3 (green). C13 position is indicated. **B,** Natural R1 congeners, dcSTX (left) and GTX5 (right). **C,** C13-aryl congener exemplars, C13-OBz (left) and C13-NBz (right). Numbers indicate aryl ring positions. Red circle highlights amide linkage in C13-NBz.

The saxiphilin (Sxph) family of high-affinity STX binding proteins from frogs and toads (anurans)^10,11,26^ are transferrin-like ∼90-120 kDa soluble proteins bearing a single toxin binding site that accommodates a range of STX congeners with affinities in the low to mid-nanomolar range^10–12,26,27^. This property supports their ability to rescue Na_V_s from block by STX and various PSTs^10,12^ and act as toxin sponges to defend against STX poisoning^17,28^. The relative ease of manipulation and host of available binding assays^29^ make Sxphs a versatile platform for investigating the energetic and structural factors that undergird molecular recognition of STX and its congeners^10–12,29^. Studies of Sxphs from the American bullfrog (*Rana catesbeiana*), *Rc*Sxph, and High Himalaya frog (*Nanorana parkeri*), *Np*Sxph, have defined the binding code that governs STX recognition by Sxphs^10,12^. This molecular recognition code comprises a set of charged, cation-π, and water-mediated interactions that coordinate the tricyclic bis-guanidinium toxin core together with hydrophobic residues that engage the carbamate moiety^10–12^. The high-affinity Sxph-toxin interaction follows a ‘lock and key’ mechanism involving minimal structural reorganization of the STX binding site upon toxin engagement^10–12^ and shares similarities with the toxin receptor site in Na_V_s^11,12^. Studies of a series of sulfated (gonyautoxins GTX2/3; GTX5, also known as B1^2^; and C1/C2) and decarbamoylated (dcSTX and dcGTX2/3) congeners^12^ have established that the Sxph binding poses of these structurally varied toxins are identical. However, this shared binding mode that engages a largely rigid binding pocket comes at the cost of reduced affinity for congeners having modifications that interfere with key interactions, such as sulfation at C11 (GTX2/3), loss of contacts in decarbamoylated toxins, and N1 hydroxylation^12^. Whether similar rules affect Sxph interactions with other PSTs and derivatives thereof is not known.

The C13 position of STX (R1 site), also denoted C17 in some numbering schemes^21,30^, (Fig. 1A) hosts chemically diverse natural variants that include decarbamoylation (ex. dcSTX)^2^ (Fig. 1B), sulfation (ex. GTX5) (Fig. 1B), and acetate (LWTX-5)^31^ and hydroxybenzoate modifications^2,32,33^. Interactions between Sxph and the STX carbamate contribute ∼1 kcal mol^-1^ to the total binding energy^10,12^. Modifications of the R1 moiety can either weaken or enhance toxin binding, as exemplified by toxins lacking the carbamate (dcSTX) or bearing a sulfocarbamoyl group (GTX5), respectively^12^. Given the rich chemical diversity at the R1 site and the differential impact of this position on binding affinity^10,12^, we wanted to investigate how other types of R1 site modifications affect Sxph binding. Hence, we synthesized a series of C13-modified toxins (Fig. 1) and examined their interactions with Sxphs. The resulting toxins spanned the smallest change relative to STX that swapped the carbamate for an acetate group to create the cyanobacterial toxin LWTX-5^31^, to changes in toxin polarity, rigidity, and steric bulk with a series of R1 benzoate and amide derivatives (Fig. 1C). Our biophysical and structural studies of *Rc*Sxph, *Rc*Sxph mutants, and *Np*Sxph reveal that the Sxph toxin binding pocket is remarkably tolerant of these R1 site structural modifications. Further, we have discovered an unexpected conformational diversity in toxin binding modes that is dependent on the R1 substituent and the local environment around *Rc*Sxph Tyr558. Together, these results expand our understanding of the principles governing Sxph binding to diverse STX congeners and present findings that are important for advancing Sxph-based anti-PSP strategies and development of new Na_V_ modulators.

### Saxiphilins bind a diverse panel of C13-modified STX congeners

Contacts between the Sxph toxin binding pocket and the carbamate group of STX and its congeners are key contributors to binding affinity^10,12^. To probe Sxph tolerance for carbamate modifications, we used a thermofluor (TF) assay^10,12,29^ in which toxin binding to Sxph manifests as a change in the apparent protein melting temperature (ΔTm). We screened a panel of eleven natural and synthetic C13-modified STX congeners against the well-characterized Sxphs *Rc*Sxph^10–12^ and *Np*Sxph^10,12^ (Figs. 2A-B and S1). This toxin series included C13-O-acetate (LWTX-5, C13-OAc)^31^, C13-O-benzoate (C13-OBz), C13-N-benzamide (C13-NBz), and various benzoate derivatives substituted with methyl, fluorine, or hydroxyl groups at the 2-, 3-, and 4-positions of the benzoate ring (Fig. 1C), namely: C13-O-2-CH_3_-benzoate (C13-O-2-CH_3_-Ph), C13-O-2-F-benzoate (C13-O-2-F-Ph), C13-O-3-CH_3_-benzoate (C13-O-3-CH_3_-Ph), C13-O-3-F-benzoate (C13-O-3-F-Ph), C13-O-2-F-benzoate (C13-O-2-F-Ph), C13-O-4-CH_3_-benzoate (C13-O-4-CH_3_-Ph), C13-O-4-F-benzoate (C13-O-4-F-Ph), C13-O-4-OH-benzoate (C13-O-4-OH-Ph), C13-O-3,4-(OH)_2_-benzoate (C13-O-3,4-(OH)_2_-Ph). Three of these, C13-OAc (LWTX-5)^31^, C13-O-4-CH_3_-benzoate (C13-O-4-CH_3_-Ph) (GC3)^32^, and C13-O-3,4-(OH)_2_-benzoate (C13-O-3,4-(OH)_2_-Ph)(GC3a)^33^, are naturally occurring toxins.

**Figure 2.**
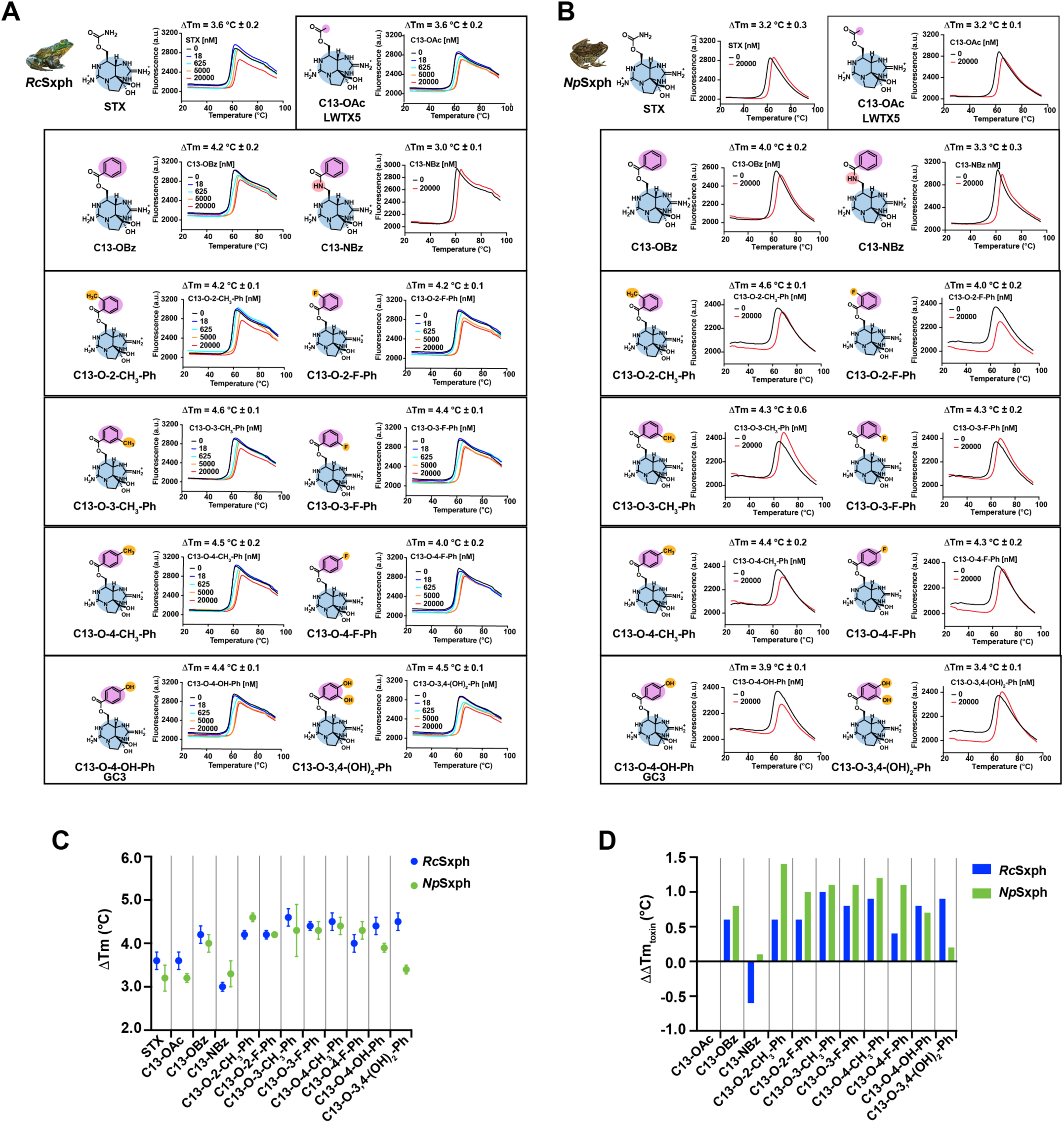
TF assays show Sxph binding preferences for C13-STX congeners. **A** and **B,** Exemplar TF assay results for **A**, *Rc*Sxph and **B,** *Np*Sxph, in the presence of 0 nM (black), 18 nM (blue), 625 nM (cyan), 5000 nM (orange), and 20,000 nM (red) of the indicated toxins. **C,** ΔTm for each toxin with *Rc*Sxph (blue) and *Np*Sxph (orange), Error bars show S.E.M. **D,** ΔΔTm_toxin_ for each toxin with *Rc*Sxph (blue) and *Np*Sxph (green). STX data for *Rc*Sxph and *Np*Sxph in ‘A’ and ‘B’ are from ^10^.

We found that *Rc*Sxph and *Np*Sxph exhibited toxin concentration-dependent increases with ΔTm values ranging from 3.0-4.6°C and 3.2-4.6°C, respectively (Figs. 2A-C, S1, and Table 1), indicating that the two homologous Sxphs can bind a diverse array of R1-substituted toxins. Most C13-modified STX derivatives displayed enhanced thermostability relative to STX (ΔΔTm_toxin_) with values that ranged from +0.1 to +1.4 °C (Fig. 2D, Table 1). Changing the STX carbamate group to acetate (C13-OAc) had no effect on ΔΔTm_toxin_ for *Rc*Sxph and *Np*Sxph (Fig. 2D, Table 1), in line with the observation the carbamate-NH_2_ group makes few interactions with the STX binding pocket in either protein^10,11^. By contrast, all aryl esters showed more pronounced ΔΔTm_toxin_ changes with both Sxphs (Fig. 2D, Table 1), suggesting that the large aryl modification is well tolerated. Replacing the C13-OBz ester linkage with an amide in C13-NBz reduced ΔΔTm_toxin_ in *Rc*Sxph (Fig. 2D, Table 1) but minimally affected ΔΔTm_toxin_ for *Np*Sxph suggesting that there are binding pocket differences between the two Sxph variants that influence C13-NBz engagement. Together, the TF data show that the C13 aryl ester and amide modified toxins are well tolerated by both Sxphs, regardless of polarity and steric bulk. These results contrast the effects of C11 sulfation^12^ and N1 hydroxylation^12^ and reflect marked differences in how the Sxph binding pocket accepts modifications at the various positions of the STX scaffold.

**Table 1.**
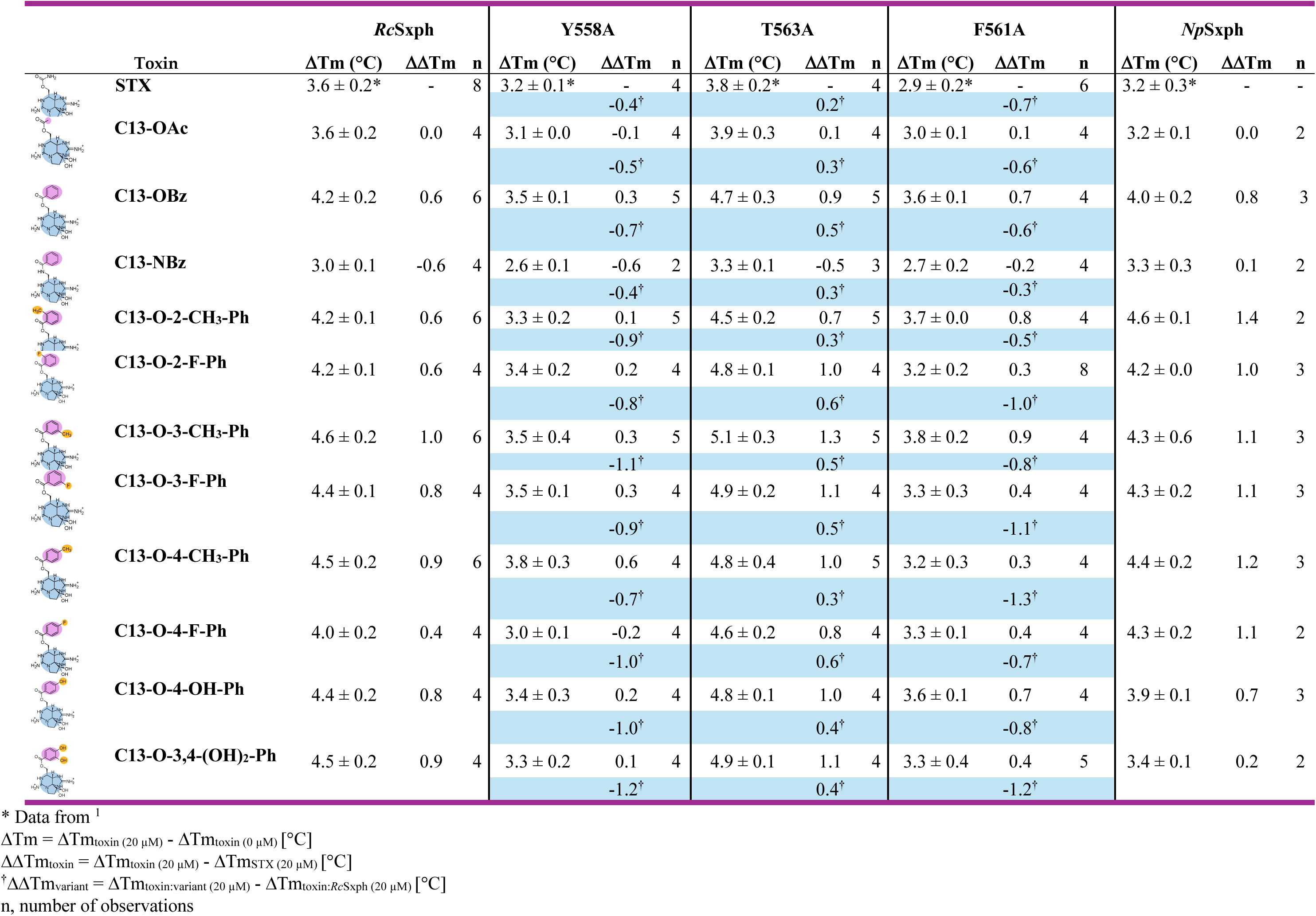
ΔTm for Sxphs in the presence of C13-modified STX congeners.

### STX binding pocket mutants indicate differential recognition of STX congeners

*Rc*Sxph Tyr558 and Phe561 form part of the toxin binding site that interacts with the STX carbamate and make important contributions to STX binding energy^10^. To probe the influence of these positions in binding to our series of C13-modified STX congeners, we used the TF assay to compare the effects of alanine substitution at *Rc*Sxph Tyr558, Phe561, and Thr563, a site adjacent to Phe561 that makes minimal contribution to STX binding energetics^10^ (Figs. 3A-E and S1, Table 1). We analyzed the TF results in two ways for each toxin, looking at changes in thermal shifts relative to STX (ΔΔTm_toxin_) (Fig. 3F, Table 1) to identify toxin specific trends and changes relative to wild-type *Rc*Sxph (ΔΔTm_variant_) to discern effects of the different alanine substitutions (Fig. 3G). None of the alanine mutations caused substantial difference in ΔΔTm_toxin_ for C13-OAc, consistent with the similarity of this toxin to STX (Fig. 3F, Table 1). By contrast, other derivatives showed differential responses to the *Rc*Sxph mutations. The abilities of C13-OBz and related aryl esters to affect ΔΔTm_toxin_ were all diminished for the Y558A mutation (Fig. 3F, Table 1). Comparing ΔΔTm_toxin_ for F561A revealed little discrimination between STX and C13-OBz, C13-O-2-CH_3_-Ph, C13-O-3-CH_3_-Ph, and C13-O-4-OH-Ph, but showed noticeably smaller ΔΔTm_toxin_ for C13-NBz, C13-O-2-F-Ph, C13-O-3-F-Ph, C13-O-4-CH_3_-Ph, and C13-O-4-(OH)_2_-Ph (Fig. 3F, Table 1). The T563A change increased ΔΔTm_toxin_ for all C13-aryl esters but decreased it for C13-NBz (Fig. 3F, Table 1), suggesting that there are differences in how C13-aryl esters and the amide compound interact with the *Rc*Sxph binding pocket.

**Figure 3.**
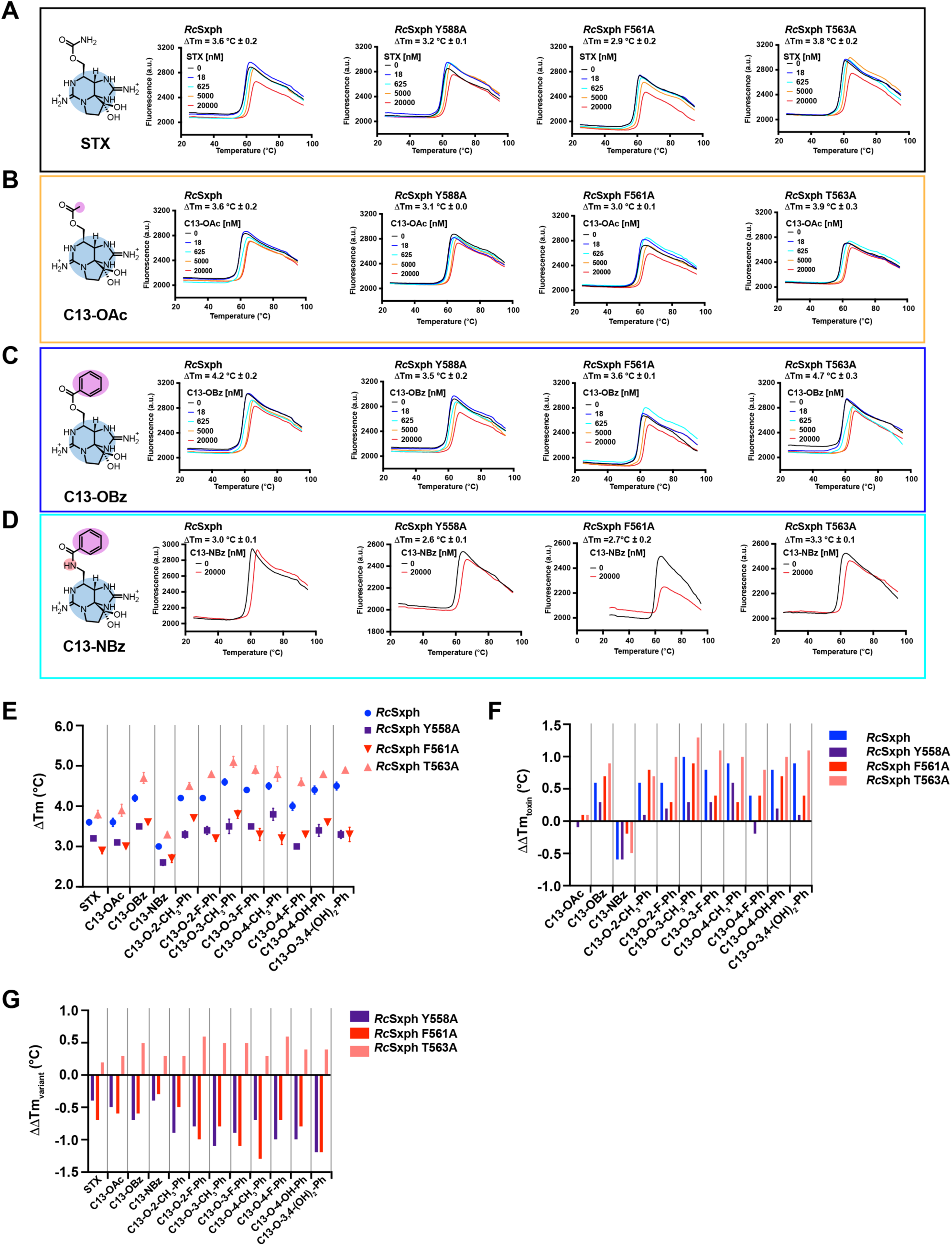
STX binding pocket mutations affect C13-STX congener binding. **A-C**, Exemplar TF assay results for *Rc*Sxph, *Rc*Sxph Y558A, *R*cSxph F561A, and *Rc*Sxph T563A in the presence of **A**, STX **B,** C13-OAc, **C,** C13-OBz, and **D,** C13-NBz in the presence of 0 nM (black), 18 nM (blue), 625 nM (cyan), 5000 nM (orange), and 20,000 nM (red) of the indicated toxins. **E,** ΔTm for each toxin with *Rc*Sxph (blue circles), *Rc*Sxph Y558A (purple squares), *Rc*Sxph F561A (red inverted triangles), and *Rc*Sxph T563A (orange triangles). Error bars show S.E.M. **F,** ΔΔTm_toxin_ for each toxin with *Rc*Sxph (blue), *Rc*Sxph Y558A (purple), *Rc*Sxph F561A (red), and *Rc*Sxph T563A (orange). **G** ΔΔTm_variant_ for each toxin with *Rc*Sxph Y558A (purple), *Rc*Sxph F561A (red), and *Rc*Sxph T563A (orange). Data in panel ‘A’ is from ^10^.

When evaluated relative to wild-type *Rc*Sxph (ΔΔTm_variant_), all three mutations had similar effects across all tested toxins. Both Y558A and F561A lowered ΔΔTm_variant_ (Fig. 3G, Table 1) indicating weaker interactions with *Rc*Sxph, although the magnitude of the changes varied (∼-0.3° to −1.3 °C). By contrast, T563A yielded a small but uniform increase of ΔΔTm_variant_ for all congeners (Fig. 3G). Collectively, analysis of the TF data suggests that all three positions (558, 561, 563) contribute to binding of C13-modified toxins, consistent with the location of these residues near the carbamate in STX-bound complexes^10,12,29^, and that Tyr558 and Phe561 are of particular importance.

C13-NBz showed lower ΔTms compared with the C13-OBz series across all Sxph variants (Figs. 2C and 3E, Table 1) and had uniformly different behavior from the C13-aryl esters relative to STX (Figs. 2D and 3F, Table 1). Although ΔTm changes generally follow alterations in binding affinity^34,35^, our previous studies revealed that for Sxphs, the correlation between ΔTm and ΔΔG for binding is more robust when changes are >1 kcal mol^-1^ ^10^. Hence, we decided to use a fluorescence polarization competition assay (FPc) ^12,36^ to measure affinities (Kds) directly and compare how STX, C13-OBz, and C13-NBz compete with fluorescein-labelled STX (F-STX) against *Rc*Sxph, *Np*Sxph, and *Rc*Sxph mutants. We focused on C13-OBz as a representative member of the aryl ester series. FPc measurements for STX binding to *Rc*Sxph Y558A, F561A, and T563A, and *Np*Sxph yielded Kd values in line with previously measured Kds using FP and isothermal titration calorimetry^10^ and confirmed the ∼10-fold destabilization to STX binding caused by the F561A change relative to wild-type *Rc*Sxph (Figs. 4A and D, Table 2). Both C13-OBz and C13-NBz showed slightly reduced affinities relative to STX for *Rc*Sxph (Kd =13.0 ± 2.9, 17.7 ± 2.3, and 8.9 ± 1.6 nM, respectively) (Figs. 4B-C, Table 2). Interestingly, Y558A enhanced binding for both toxins relative to wild-type *Rc*Sxph (ΔΔG_variant_ ∼-0.5 to −0.7 kcal mol^-1^) (Fig. 4D), similar to how this mutation affects STX binding^10^. Notably, *Np*Sxph, a Sxph that bears an isoleucine at the position equivalent to Tyr558 (*Np*Sxph Ile559), exhibited nearly identical increases in affinity (ΔΔG_variant_ ∼-0.6 to −0.7 kcal mol^-1^) for both C13-OBz and C13-NBz compared to *R*cSxph (Fig. 4D, Table 2).

**Figure 4.**
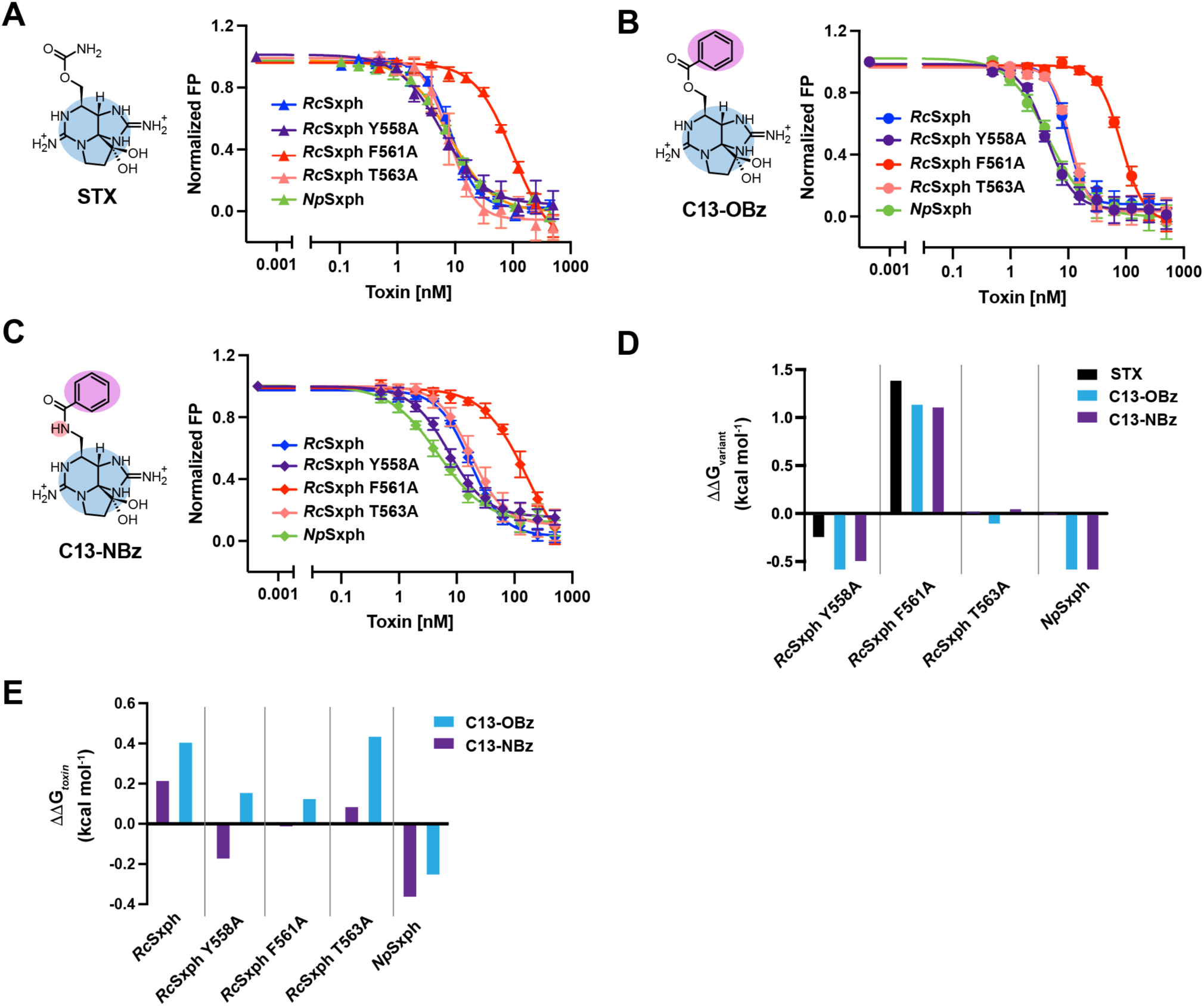
FPc studies of select C13-STX congeners. **A-C**, Exemplar FP competition assays, for **A,** STX, **B,** C13-OBz, and **C,** C13-NBz with *Rc*Sxph (blue), *Rc*Sxph Y558A (purple), *Rc*Sxph F561A (red), *Rc*Sxph T563A (orange), and *Np*Sxph (green). **D,** ΔΔG_variant_ for each Sxph with STX (black), C13-OBz (blue), and C13-NBz (purple). **E,** ΔΔG_toxin_ for each Sxph with C13-OBz (blue), and C13-NBz (purple). Error bars show SD. **A-C**, shows toxin structures colored as in Fig. 1.

**Table 2.**
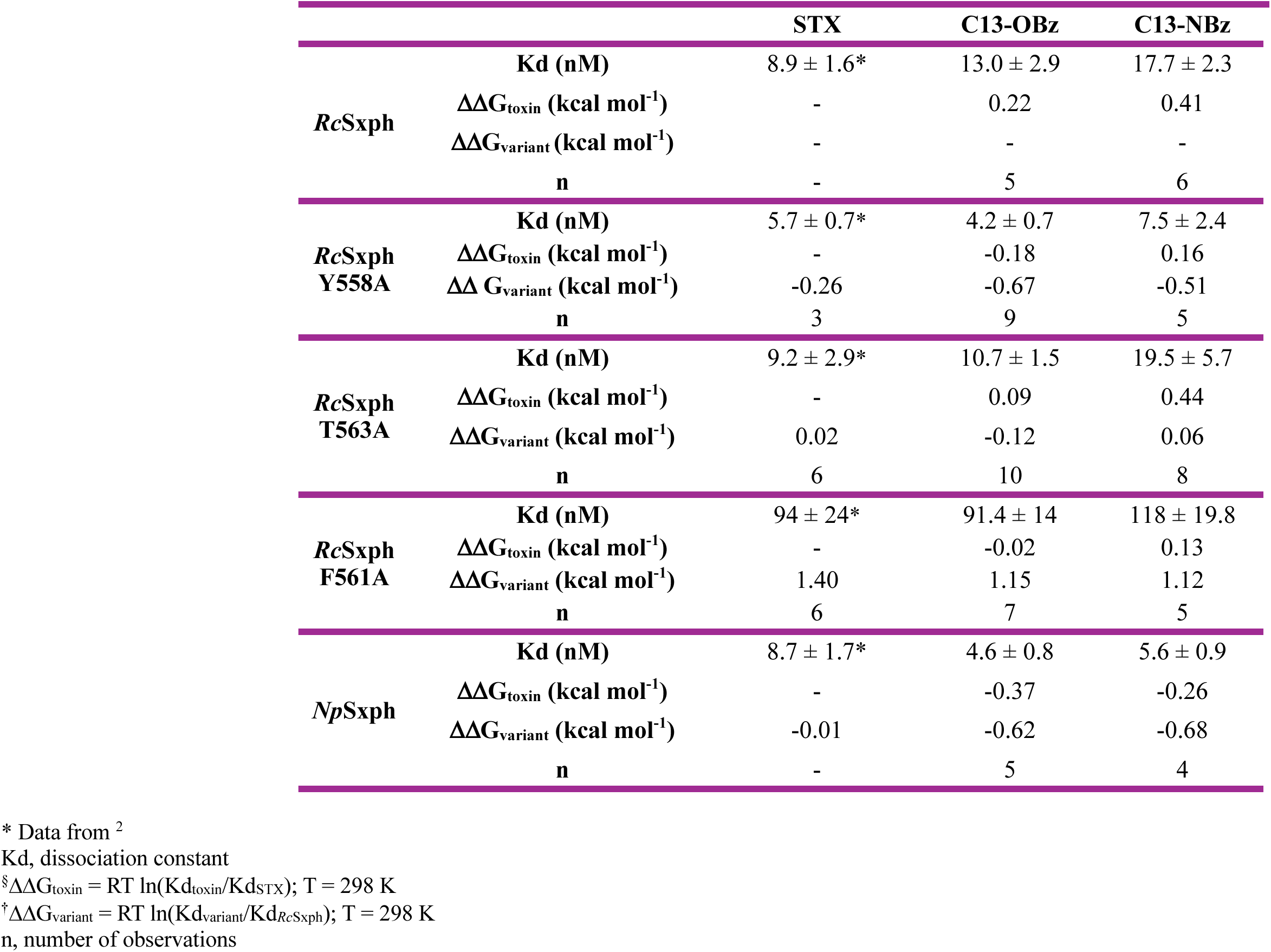
Fluorescence polarization competition (FPc) data comparison.

The increased binding affinity of *Rc*Sxph Y558A and *Np*Sxph for C13-OBz and C13-NBz agrees with previous work demonstrating that both Sxphs have enhanced STX binding originating from relief of a steric clash at the *Rc*Sxph Tyr558 position with the STX core^10^ and suggest an important role for this site in C13-aryl congener binding. Contrasting the effects of Y558A, the F561A mutation reduced binding of all three toxins (ΔΔG_variant_ ∼ 1.1-1.4 kcal mol^-1^) (Fig. 4D, Table 2) recapitulating the effect of this mutation on STX^10^. Finally, the FPc assays indicated that the T563A change was essentially neutral across all tested toxins with no energy penalty for accommodating diverse modifications compared to wild-type *Rc*Sxph (Fig. 4D, Table 2). Together, these data reveal that despite chemical and structural differences of C13 modifications, the relative energetic contributions of Tyr558, Phe561, and Thr563 follow their effects on STX.

Analysis of the changes relative to STX (ΔΔG_toxin_) shows that both C13-OBz and C13-NBz have slightly lower affinity than STX for *Rc*Sxph and T563A (ΔΔG_toxin_ ∼0.1-0.4 kcal mol^-1^) (Fig. 4E, Table 2). The Y558A and F561A mutations diminish these differences between STX and the C13-aryl ester derivatives (Fig. 4E, Table 2). Notably, this trend is reversed for *Np*Sxph, where C13-OBz and C13-NBz display comparable affinities that are both more favorable than STX (ΔΔG_toxin_ ∼ −0.3 to −0.4 kcal mol^-1^) (Fig. 4E, Table 2). These data further support the conclusions from the TF studies that highlight key differences in the binding interactions of this family of toxins with *Rc*Sxph and *Np*Sxph.

### Structures of Sxph:STX congener complexes reveal two toxin binding conformations

To understand the molecular details of how the C13 toxin derivatives interact with Sxph, we determined X-ray crystal structures of C13-OBz complexes with *Rc*Sxph (2.40Å), *Np*Sxph (1.90Å), *Rc*Sxph Y558A (2.45Å), *Rc*Sxph F561A (2.40Å), and *Rc*Sxph T563A (2.45Å), an *Np*Sxph:C13-NBz complex (1.95Å), an *Rc*Sxph-F561A:STX complex (2.35Å), and apo-structures of *Rc*Sxph F561A (2.23Å) and *Rc*Sxph T563A (2.50Å) (Fig. S2, Table S1). As with previous *Rc*Sxph structures, we used chain B for analysis as this chain was better defined in the asymmetric unit^10,11^. All complexes show toxin bound to the STX binding site with a 1:1 toxin:Sxph stoichiometry identical to previously determined Sxph:STX and Sxph:STX congener complexes^10–12^ (Figs. 5 and S2-S4). There are no large conformational changes in the Sxph core (N- and C-lobes) and thyroglobulin domains (Thy1-1 and Thy1-2) induced by toxin binding (Fig. S2)^10–12^ (root mean square deviation of Cα positions (RMSD_Cα_) = 0.507, 0.397, 0.257, 0.466, 0.496, 0.202, and 0.203 Å for *Rc*Sxph:C13-OBz, *Rc*Sxph-Y558A:C13-OBz, *Rc*Sxph-F561A:C13-OBz, *Rc*Sxph-F561A:STX, *Rc*Sxph-T563A:C13-OBz, *Np*Sxph:C13-OBz, and *Np*Sxph:C13-NBz, respectively).

**Figure 5.**
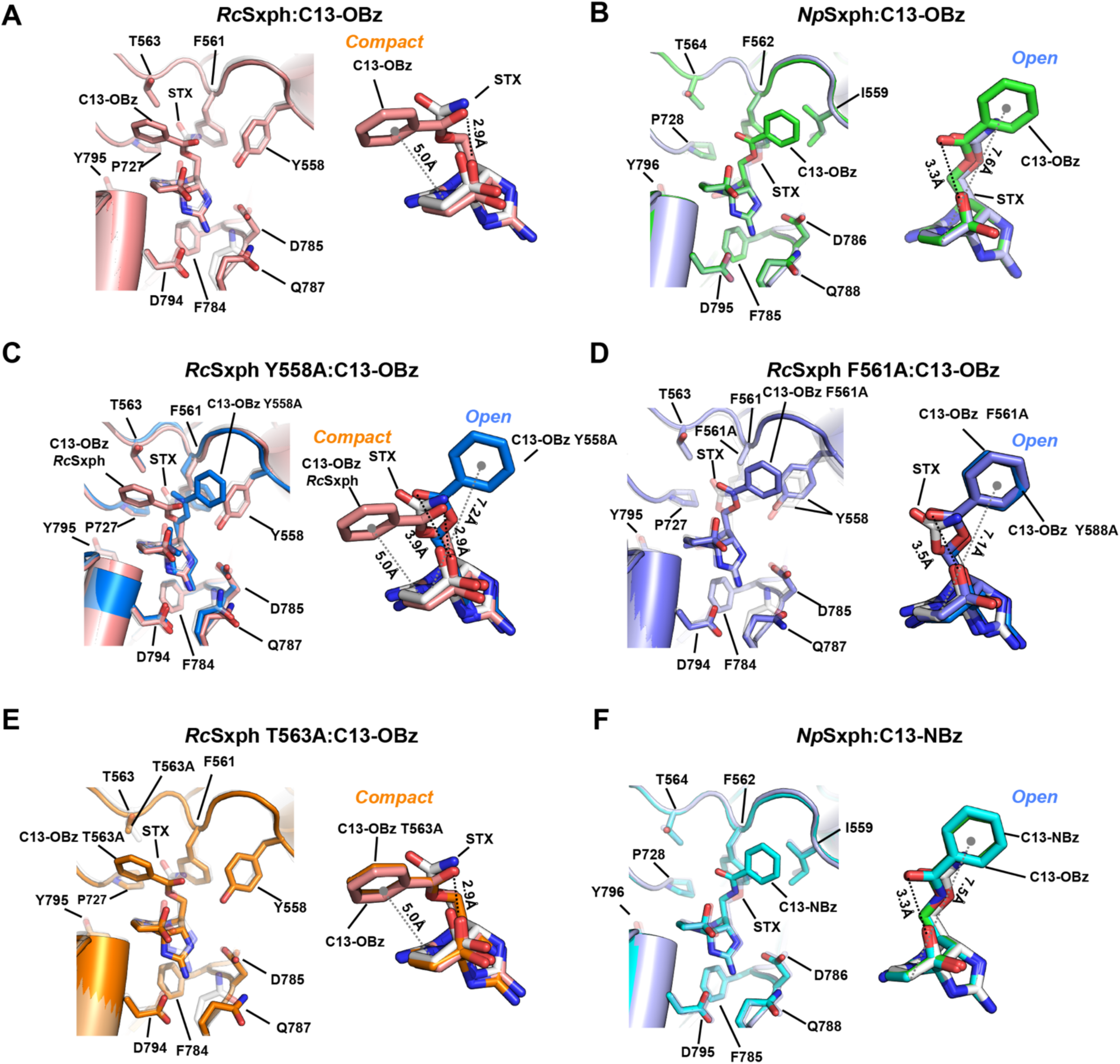
Comparison of C13-STX congener and STX binding modes. Panels depict binding pocket comparisons (left) and bound toxin conformations (right). **A,** *Rc*Sxph:C13-OBz (salmon) and *Rc*Sxph:STX (6O0F) (white)^11^. **B,** *Np*Sxph:C13-OBz (green) and *Np*Sxph:STX (light blue) (8D6M)^10^. **C,** *Rc*Sxph 558A:C13-OBz (marine), *Rc*Sxph:C13-OBz (salmon), and *Rc*Sxph:STX (6O0F) (white)^11^. **D,** *Rc*Sxph F561A:C13-OBz (slate) and *Rc*Sxph:STX (6O0F) (white)^11^. **E,** *Rc*Sxph T563A:C13-OBz (orange), and *Rc*Sxph:STX (6O0F) (white)^11^. Right panel also shows C13-OBz from the *Rc*Sxph:C13-OBz complex (salmon). **F,** *Np*Sxph:C13-NBz (cyan), and *Np*Sxph:STX (light blue) (8D6M)^10^. C13-OBz, C13-NBz, and STX are shown as sticks.

All *Rc*Sxph and *Np*Sxph complexes with C13 toxin derivatives show binding pocket residue interactions with the tricyclic toxin core that are identical to that observed for STX (Figs. 5A-F and S4)^10,11^, dcSTX^12^, and various sulfated STX congeners^12^. *Rc*Sxph Glu540/*Np*Sxph Glu541 coordinate the STX six membered guanidinium ring. *Rc*Sxph Asp785 and Asp794/*Np*Sxph Asp786 and Asp795 coordinate the five membered guanidinium ring, and *Rc*Sxph Phe784/*Np*Sxph Phe785 engage the STX five membered guanidinium ring via a cation-π interaction at the base of the binding pocket (Fig. S4). The STX binding site residues display minimal changes relative to the apo forms as observed in prior Sxph structures^10,11^, except for the characteristic inward movement of *Rc*Sxph Asp794/*Np*Sxph Asp795 upon toxin binding (Figs. S3B-D). Hence, the ‘lock and key’ mode of binding observed for STX^10,11^ and a panel of decarbamoylated and sulfated congeners^12^ is preserved for these C13-aryl ester and amide variants.

Despite these canonical interactions with the STX core, the C13-OBz complexes revealed an unexpected variation in toxin binding modes of the aryl ester group. Depending on the structure, we found one of two distinct conformations, denoted as ‘compact’ and ‘open’ (Figs. 5A-F). The *Rc*Sxph:C13-OBz complex showed the ‘compact’ ligand pose in which the carbonyl oxygen and center of the phenyl ring are close to the C12 β-hydroxyl group and the C11 position of the pyrrole ring at distances of 2.9Å and 5.0Å, respectively. In this pose, the phenyl ring contacts the part of the binding pocket made by Thr563 (Fig. 5A). By contrast, the same toxin bound to *Np*Sxph displays the ‘open’ conformer in which the C13-OBz phenyl ring is flipped to the opposite side of the binding pocket and contacts Ile559 (Figs. 5B and S4B). In this ‘open’ conformation the distance between the phenyl ring and both the C12 β-hydroxyl group and C11 center is much larger than in the ‘compact’ pose (3.3Å and 7.6Å, respectively) (Fig. 5B). Strikingly, the presence of the ‘open’ or ‘compact’ conformer correlates with whether a C13-derivative is a better binder than STX (ΔΔG_toxin_ < 0), as in *Np*Sxph, or worse binder than STX (ΔΔG_toxin_ > 0), as in *Rc*Sxph (Fig. 4E, Table 2), respectively.

In line with this observation, the *Rc*Sxph Y558A:C13-OBz complex (Figs. 5C and S4C), a mutant to which C13-OBz binds more strongly than STX (Fig. 4E, Table 2), also shows the ‘open’ conformer. The ‘open’ conformer is also seen in the *Rc*Sxph F561A:C13-OBz complex (Figs. 5D and S4E), a mutant for which there is minimal energetic difference between STX and C13-OBz (Fig. 4E, Table 2). Interestingly, in the apo*-Rc*Sxph F561A structure, the Tyr558 residue sits ∼2Å away from its apo-*Rc*Sxph binding pocket position^11^ (Fig. S3A). In the *Rc*Sxph F561A:C13-OBz, the conformations of Tyr558 and its loop match the apo-*Rc*Sxph F561A structure (Figs. 4D and S3A-B), indicating that the Tyr558 site is influenced by the Phe561. Examination of the *Rc*Sxph F561A:STX complex revealed a repositioning of Tyr558 and its loop relative to both the apo-*Rc*Sxph F561A structure (Fig. S3B) and the *Rc*Sxph F561A:C13-OBz complex (Fig. S3C). Hence, the loss of the Phe561 sidechain enables Tyr558 mobility.

Contrasting the effects of the Y558A and F561A mutants, structural determination of the apo- and C13-OBz bound *Rc*Sxph T563A (Figs. 5E and S3D) showed the compact C13-OBz conformation, aligning with the ΔΔG_toxin_ trend (Fig. 4E, Table 2). Apart from the standard movements of Tyr558 and Asp785^10–12^, the *Rc*Sxph T563A structures display few toxin binding pocket changes between the apo- and C13-OBz bound forms (Fig. 5E and S3D). Overall, this set of C13 aryl ester bound structures demonstrates that relief of clashes with *Rc*Sxph Tyr558 by replacement with a smaller residue (ex. *Rc*Sxph Y558A or *Np*Sxph where the equivalent position is Ile559) or its movement away from the STX binding site (*Rc*Sxph F561A) yields the ‘open’ toxin conformer and emphasize the important role of this position in influencing toxin binding affinity^10–12^.

Consistent with the idea that the *Rc*Sxph Tyr558 position is a key determinant for the binding of the ‘open’ versus ‘compact’ form of the C13-aryl ester congeners, the *Np*Sxph:C13-NBz structure also exhibits an ‘open’ ligand conformation (Fig. 5F). These findings establish that the C13-aryl ester and C13-NBz congeners can adopt this pose regardless of the ester or amide linkage and supports the idea that the ‘open’ conformer is correlated with tighter binding (Figs. 4D-E, Table 2). Together, these structures reveal that C13-aryl ester STX derivatives can adopt two distinct binding conformers governed by local steric constraints near the C13 group that can make stable binding interactions with the toxin binding site.

### Bound waters match those found in other STX congener complexes

A set of water-mediated Sxph:toxin interactions has been shown to be important for Sxph binding to STX and various congeners^12^. The common binding mode among all Sxph:toxin structures with the STX tricylic core combined with the variability in the toxin conformations at the carbamate position prompted us to examine whether there are any changes in the water network in the C13-OBz and C13-NBz complexes. As with prior studies^10,12^, the *Np*Sxph structures are higher resolution than the *Rc*Sxph complexes and allow for better analysis of bound waters. Comparison of the *Np*Sxph:C13-OBz and *Np*Sxph:C13-NBz identified the same water-mediated networks identified previously^12^. These include Wat541 that bridges the Glu541 sidechain with the Glu784 backbone carbonyl (Fig. S3E), Wat786 found between the Asp786 sidechain and the STX N7 atom (Fig. S3F), and Wat795 that connects with the STX 14-OH and N9 positions (Fig. S3F). The conserved nature of this water-mediated network together with the canonical direct sidechain interactions provide clear evidence that Sxphs bind the STX tricyclic core in a manner that is independent of the identity of R1 moiety^11,12^.

## Discussion

The diverse modifications to the STX scaffold at three principal sites (R1, R2, and R3) (Fig. 1A) strongly influence the binding affinities and toxicities within this family of small molecule bis-guanidinium poisons^1,2,6^. Establishing the rules by which these structural variations shape toxin-target interactions is an essential goal for developing countermeasures against PSTs^17^ and for creating STX-inspired molecules to probe and modulate Na_V_ function, especially in context of isoform selective agents^21,24,25^.

Studies of Sxphs, soluble high affinity STX binding proteins from frogs and toads (anurans), have defined basic principles governing recognition of many PSTs^10–12^. These data show that high-affinity binding arises from a largely pre-organized binding pocket that undergoes minimal conformational change upon toxin engagement. Remarkably, this ‘lock and key’ binding mechanism shares core STX recognition principles with Na_V_s^11,12^. However, this structural rigidity comes at a cost of reduced affinity for certain STX congeners, particularly those bearing R2 hydroxylation or R3 sulfation^12^. Given these constraints, whether and to what extent the Sxph toxin binding site can adapt to accommodate the chemical diversity in natural and synthetic PSTs remains an unresolved issue.

Structural studies of *Rc*Sxph^10,11^ and *Np*Sxph^10,12^ have shown that STX and its congeners bind in a conserved pose in which the tricyclic core occupies the deepest point of the binding pocket while the R1 carbamate extends towards the aqueous opening. This orientation tolerates carbamate modifications such as sulfation (GTX5)^12^ and the addition of a six carbon-linked fluorescein dye to STX (F-STX)^10^ without diminishing affinity^10,12^. Nevertheless, the carbamate group is important for binding affinity, as decarbamoylation (dcSTX) or mutation of the carbamate-interacting residues reduces toxin affinity by ∼0.5-1.0 kcal mol^-1^ ^10,12^.

Here, our studies of C13-modified toxins unmask a previously unappreciated plasticity in the interaction between the Sxph toxin binding site and toxin. We find that the largely rigid Sxph toxin binding pocket can accommodate diverse C13 substitutions and that natural and designed local changes can tune toxin binding affinity and influence toxin binding conformation. This structural flexibility of the bound toxin was unexpected. We observed two distinct binding conformations for the C13-aryl esters, termed ‘compact’ and ‘open’ (Fig. 6A) that are dictated by the local architecture of the binding pocket near *Rc*Sxph Tyr558 (Fig. 6B-C). Relief of a steric clash with Tyr558 is the critical determinant that favors binding of the ‘open’ form (Fig. 6C) as seen in the structures of *Rc*Sxph Y558A, *Rc*Sxph F561A, and *Np*Sxph having isoleucine at the Tyr558 equivalent position. These changes that favor the ‘open’ form enhance toxin binding (Fig. 4E). Hence, the ‘compact’ conformation appears to be energetically strained. Contrasting this conformational plasticity at the R1 position, the protein-ligand contacts and structured water network that stabilize the tricyclic STX core are preserved and serve as the anchor that enables the conformational variability at the R1 position. Thus, different portions of the toxin scaffold are recognized through separate modes. The identification of two binding poses for C13-aryl esters of STX to Sxphs has implications for toxin recognition by other STX targets, particularly Na_V_s, where local changes in the toxin binding site may similarly influence toxin conformational preferences and provide a basis for isoform-specific interactions^19^.

**Figure 6.**
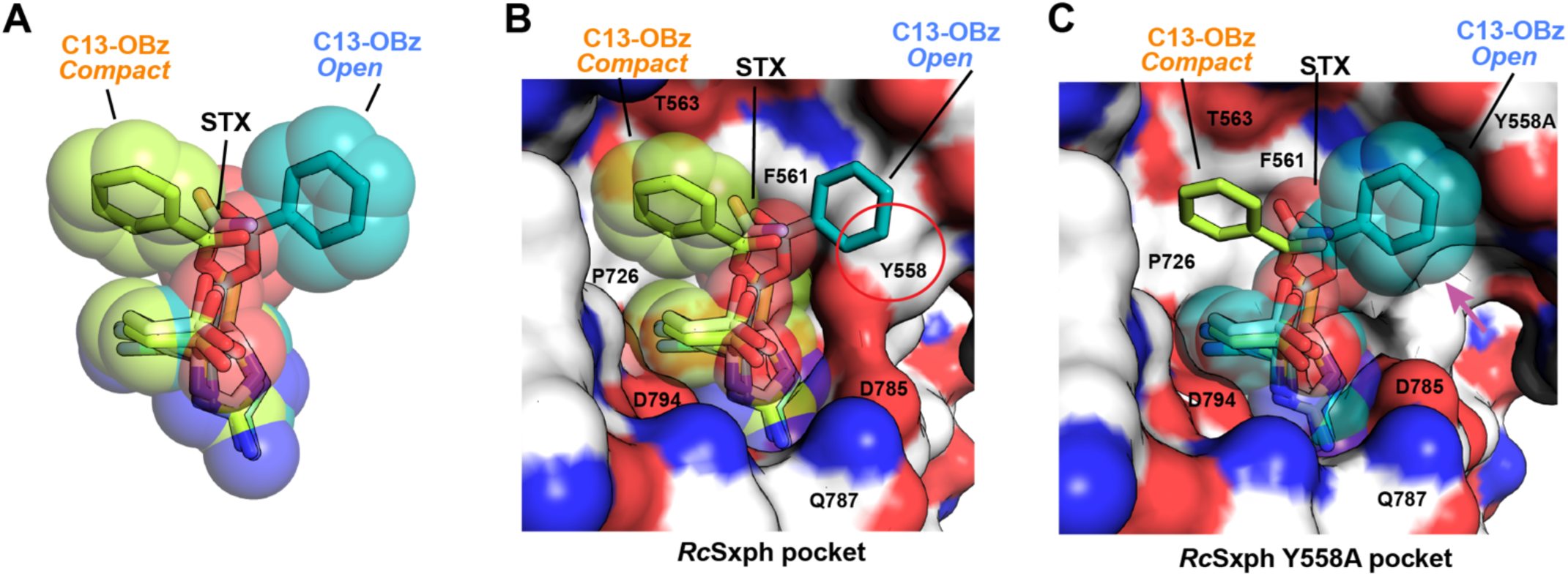
Conformational variability of aryl ester derivatives is influenced by the *Rc*Sxph Tyr558 position. **A**, Space filling models of compact (lime) and open (teal) forms of C13-OBz compared with STX (white). **B,** and **C,** Comparison of compact and open forms for **B,** the *Rc*Sxph pocket and **C,** the *Rc*Sxph Y558A **pocket.** Red circle indicates the open form clash with Tyr558. Magenta arrow indicates space opened by the Y558A mutation.

Our data show that Sxphs can effectively bind a broad range of natural and synthetic toxins and elucidate how the ‘lock and key’ receptor-ligand binding model can retain sufficient local plasticity to engage structurally diverse ligands. These qualities, together with prior studies showing binding for a variety of sulfated toxins^12^, suggest that the Sxph family can be developed as broadly acting PSTs sensors or counteragents. Exploiting natural variations in the residues comprising the Sxph toxin binding pockets from different frogs and toads^10^ may offer a way to capitalize on the balance between the ‘lock and key’ interaction mode and binding pocket plasticity to create Sxphs or Sxph derivatives capable of binding diverse toxin congeners with high affinity. Such factors will be important for developing Sxph-inspired toxin sponge molecules that can broadly counteract PSTs ^1,2,6^. Together, our findings highlight an unexpected interplay between toxin and binding pocket adaptability, demonstrate that two, stable toxin binding modes can occur within highly similar, rigid binding sites, and provide key information for the design of new STX-based probes and agents to study Na_V_s.

## Materials and Methods

### Sxph production and purification

*Rana catesbeiana* saxiphilin (*Rc*Sxph) (GenBank: U05246.1)^11^, *Rc*Sxph mutants^10^ and *Nanorana parkeri* saxiphilin (*Np*Sxph) (GenBank: XM_018555331.1)^10^ were produced by expression as secreted proteins from insect cells using baculoviruses and were purified as previously described^10,11,29^. In brief, Sxphs carrying a C-terminal 3C protease cleavage site, green fluorescent protein (GFP), and a His_10_ tag in series were expressed in *Spodoptera frugiperda* (*Sf9*) cells using a baculovirus expression system^29^. After addition of P2/P3 baculovirus to *Sf9* cells at the dilution ratio of 1:50 (v/v), cells were incubated in a non-humidified New Brunswick Innova 44 incubator (Eppendorf, cat. no. M1282-0010) at 27°C, shaking at 130 rpm for 72h. Cells were harvested by centrifugation (JLA-8.1000 rotor, 4000*g*). The supernatant was adjusted to pH 8.0 with a final concentration of 50 mM Tris-HCl and treated with 1 mM NiCl_2_ and 5 mM CaCl_2_ to precipitate contaminants. Precipitants were removed by centrifugation (JLA-8.1000 rotor, 6200*g*), and the clarified supernatant was incubated with antiGFP nanobody-conjugated Sepharose resin for 5 hours at room temperature (23 ± 2°C). The resin was washed with 20 column volumes of a wash buffer containing 300 mM NaCl and 30 mM Tris-HCl (pH 7.4). After purification with antiGFP nanobody resin^37^, protein samples were treated with 3C protease^37^(0.2 mg mL^-1^ in the wash buffer) overnight at 4°C to remove the GFP-His tag. The cleaved eluates were collected and purified by size exclusion chromatography (SEC) using a Superdex 200 10/300 GL column (Cytiva). For the TF and FPc assays Sxphs were purified using a final SEC step in 150 mM NaCl, 10 mM HEPES (pH 7.4). Protein concentrations were determined by measuring UV absorbance at 280 nm using the following extinction coefficients calculated using the ExPASY server (https://web.expasy.org/protparam/): *Rc*Sxph, *Rc*Sxph T563A, and *Rc*Sxph F561A mutants, 96,365 M^−1^ cm^−1^; *Rc*Sxph Y558A mutant, 94,875 M^−1^ cm^−1^; *Np*Sxph, 108,980 M^−1^ cm^−1^.

### Toxin synthesis and preparation

Saxitoxin (STX) and fluorescein-labeled saxitoxin (F-STX) were synthesized, purified, and validated as outlined in refs. ^10,18,29,38^. STX and its congeners powders were directly dissolved with MilliQ water to make 1 mM stocks. F-STX powder was directly dissolved with MilliQ water to make 1 µM stock.

All reagents were obtained commercially unless otherwise noted. Organic solutions were concentrated under reduced pressure (ca. 60 Torr) by rotary evaporation. HPLC-grade CH_3_CN was obtained from commercial suppliers and used as is. N,N-Dimethylformamide (DMF) was passed through two columns of activated alumina prior to use.

Semi-preparative high-performance liquid chromatography (HPLC) was performed on a Varian ProStar model 210. Thin-layer chromatography was performed on EM Science silica gel 60 F_254_ plates (250 µm). Visualization of the developed chromatogram was accomplished by fluorescence quenching.

High-resolution mass spectra were obtained from the Vincent Coates Foundation Mass Spectrometry Laboratory at Stanford University. Samples were analyzed by LC-flow injection Electrospray Ionization/Mass Spectrometry (ESI/MS) on the Waters Acquity H-Class Plus UPLC and Thermo Exploris 240 BioPharma Orbitrap mass spectrometer scanning m/z 100–1000 Da. Acetonitrile with 0.1% formic acid at a flow rate of 0.2mL/min was used to transport the injected sample to the source directly, without chromatographic separation.

Saxitoxin derivatives were quantified by ^1^H nuclear magnetic resonance (NMR) spectroscopy on a Varian Inova 600 MHz NMR instrument using distilled DMF as an internal standard. A relaxation delay (d_1_) of 20 s and an acquisition time (at) of 10 s were used for spectral acquisition. The concentration of the toxin derivative was determined by integrating ^1^H signals corresponding to the toxin and a fixed concentration of the DMF standard.

The intermediate alcohol can be obtained through a previously described route ^39^.

Synthesis of the acetate ester (***C13-OAc***) was previously reported^19^. The benzamide (***C13-NBz***) was synthesized as described in ^21^.

### General Procedure for the Synthesis Aryl Esters (adapted from ^40^)

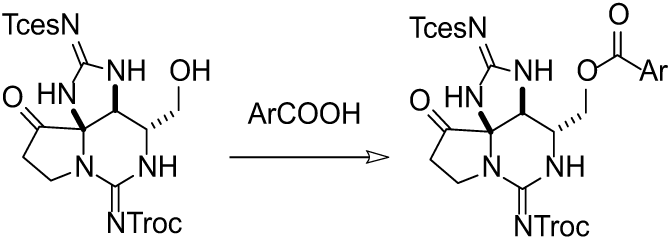

A stock solution of 80 µL of *N*-methylimidazole in 6.0 mL of CH_3_CN was freshly prepared. A 10 mL round bottom flask was charged with carboxylic acid (580 μmol, 1.2 equiv) and the alcohol starting material (30 mg, 480 μmol). A solution of *N*-methylimidazole (0.6 mL, 0.1 mmol, 2.1 equiv) was added dropwise followed by a single portion of solid *N*,*N*,*N’*,*N’*-tetramethylchloroformamidinium hexafluorophosphate (TCFH, 16 mg, 580 μmol, 1.2 equiv). The reaction progress was monitored by thin-layer chromatography and the contents were stirred until the starting alcohol had been largely consumed (24–48 h). Following this time, the reaction mixture was transferred to a separatory funnel with 20 mL of *i*-PrOAc and 20 mL of H_2_O. The organic layer was collected, dried over Na_2_SO_4_, filtered, and concentrated under reduced pressure to a white solid. Purification of this material by preparatory thin-layer chromatography (70% hexanes/EtOAc) yielded the desired aryl ester as a white solid.

### General Procedure for the Deprotection of STX-esters (adapted from ^41^)

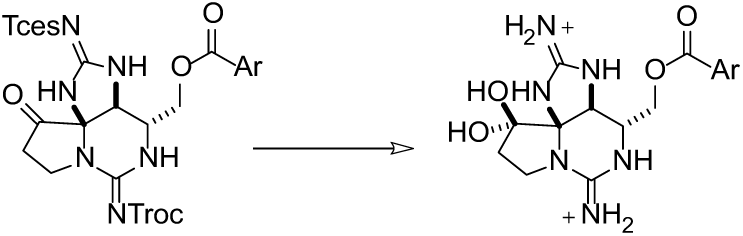

To a solution of aryl ester (18.0 μmol) in 4.0 mL of 3:1 MeOH/H_2_O was added 180 μL of acetic acid. The mixture was stirred for 30 min then PdCl_2_ (1.6 mg, 0.5 equiv) was added. The suspension was sparged with N_2_ for 5 min and then with H_2_ for 5 min. The flask was fitted with a balloon of H_2_ (1 atm) and the contents were stirred for 3 h. Following this time, the mixture was filtered through a 0.2 µm polytetrafluoroethylene (PTFE) syringe filter. The flask and filter were rinsed with 4 mL of MeOH and 4 mL of a 0.5 N aqueous AcOH solution, and the combined filtrates were concentrated under reduced pressure to a thin-film residue. This material was dissolved in 2 mL 0.5 N aqueous AcOH and the mixture was stirred for 1 h, then frozen and lyophilized to remove all volatiles. The yellow residue was purified by reversed-phase HPLC (0–50% CH_3_CN in H_2_O/heptafluorobutyric acid). Fractions were collected, frozen, and lyophilized to remove all volatiles. Fractions were quantified by ^1^H NMR integration and lyophilized again to yield the desired bis-guanidinium aryl ester.

### C13-OBz

Benzoate: Synthesized according to the above general procedure to yield the title product (7%).

**^1^H NMR** (600 MHz, D_2_O) δ 8.06 (d, 2H), 7.72 (t, 1H), 7.57 (t, J = 7.8 Hz, 2H), 4.87 (s, 1H), 4.69–4.63 (m, 1H), 4.39–4.33 (m, 1H), 4.07–4.01 (m, 1H), 3.85–3.78 (m, 1H), 3.60–3.53 (m, 1H), 2.45–2.39 (m, 1H), 2.39– 2.30 (m, 1H) ppm

**HRMS** (ESI^+^) calcd for C_16_H_20_N_6_O_4_ 360.1546 found 361.1617 (MH^+^)

### C13-O-2-CH_3_-Ph

2-Methylbenzoate: Synthesized according to the above general procedure to yield the title product (10%).

**^1^H NMR** (600 MHz, D_2_O) δ 7.93 (d, 1H), 7.57 (t, J = 1.4 Hz, 1H), 7.44–7.34 (m, 2H), 4.89 (s, 1H), 4.66– 4.59 (m, 1H), 4.41–4.34 (m, 1H), 4.08–4.00 (m, 1H), 3.85–3.78 (m, 1H), 3.60–3.49 (m, 1H), 2.56 (s, 3H), 2.48–2.40 (m, 1H), 2.40–2.31 (m, 1H) ppm

**HRMS** (ESI^+^) calcd for C_17_H_22_N_6_O_4_ 374.1703 found 375.1772 (MH^+^)

### C13-O-2-F-Ph synthesis

2-Fluorobenzoate: Synthesized according to the above general procedure to yield the title product (42%).

**^1^H NMR** (600 MHz, D_2_O) δ 7.99 (t, J = 1.8 Hz, 1H), 7.77–7.70 (m, 1H), 7.36 (t, J = 7.6 Hz, 1H), 7.34–7.27 (m, 1H), 4.89 (s, 1H), 4.70–4.63 (m, 1H), 4.43–4.35 (m, 1H), 4.08–4.01 (m, 1H), 3.86–3.79 (m, 1H), 3.61– 3.51 (m, 1H), 2.47–2.40 (m, 1H), 2.40–2.33 (m, 1H) ppm

**HRMS** (ESI^+^) calcd for C_16_H_19_FN_6_O_4_ 378.1452 found 379.1525 (MH^+^)

### C13-O-3-CH_3_-Ph

3-Methylbenzoate: Synthesized according to the above general procedure to yield the title product (8.5%).

**^1^H NMR** (600 MHz, D_2_O) δ 7.91 (s, 1H), 7.88 (d, J = 7.8 Hz, 1H), 7.58 (d, J = 7.6 Hz, 1H), 7.48 (t, J = 7.7 Hz, 1H), 4.89 (s, 1H), 4.70–4.63 (m, 1H), 4.39–4.32 (m, 1H), 4.08–4.01 (m, 1H), 3.87–3.79 (m, 1H), 3.62– 3.55 (m, 1H), 2.46–2.41 (m, 4H), 2.40–2.33 (m, 1H) ppm

**HRMS** (ESI^+^) calcd for C_17_H_22_N_6_O_4_ 374.1703 found 375.1772 (MH^+^)

### C13-O-3-F-Ph synthesis

3-Fluorobenzoate: Synthesized according to the above general procedure to yield the title product (32%).

**^1^H NMR** (600 MHz, D_2_O) δ 7.89 (d, J = 7.8 Hz, 1H), 7.79 (d, J = 2.2 Hz, 1H), 7.62–7.56 (m, 1H), 7.51– 7.45 (m, 1H), 4.89 (s, 1H), 4.71–4.62 (m, 1H), 4.44–4.32 (m, 1H), 4.10–4.00 (m, 1H), 3.90–3.78 (m, 1H), 3.65–3.53 (m, 1H), 2.47–2.41 (m, 1H), 2.41–2.33 (m, 1H) ppm

**HRMS** (ESI^+^) calcd for C_16_H_19_FN_6_O_4_ 378.1452 found 379.1523 (MH^+^)

### C13-O-4-F-Ph synthesis

4-Fluorobenzoate: Synthesized according to the above general procedure to yield the title product (36%).

**^1^H NMR** (600 MHz, D_2_O) δ 8.16–8.04 (m, 2H), 7.28 (t, 2H), 4.87 (s, 1H), 4.68–4.58 (m, 1H), 4.39–4.30 (m, 1H), 4.07–4.00 (m, 1H), 3.89–3.77 (m, 1H), 3.61–3.52 (m, 1H), 2.46–2.40 (m, 1H), 2.40–2.32 (m, 1H) ppm

**HRMS** (ESI^+^) calcd for C_16_H_19_FN_6_O_4_ 378.1452 found 379.1523 (MH^+^)

### C13-O-4-CH_3_-Ph

4-Methylbenzoate: Synthesized according to the above general procedure to yield the title product (22%).

**^1^H NMR** (400 MHz, CD_3_OD) δ 7.89 (d, J = 8.0 Hz, 2H), 7.24 (d, J = 8.2 Hz, 2H), 4.75 (s, 0H), 4.53–4.38 (m, 1H), 4.29–4.18 (m, 1H), 3.90–3.80 (m, 1H), 3.80–3.70 (m, 1H), 3.52–3.38 (m, 1H), 2.35 (s, 3H), 2.32– 2.24 (m, 2H) ppm (note: C5 proton obscured by CH_3_OH solvent peak)

**HRMS** (ESI^+^) calcd for C_17_H_22_N_6_O_4_ 374.1703 found 375.1774 (MH^+^)

### C13-O-4-OH-Ph

4-Hydroxybenzoate: Synthesized according to the above general procedure to yield the title product (16%).

**^1^H NMR** (600 MHz, D_2_O) δ 7.99 (d, 2H), 7.01 (d, 2H), 4.68–4.60 (m, 1H), 4.39–4.27 (m, 1H), 4.06–3.99 (m, 1H), 3.88–3.79 (m, 1H), 3.65–3.54 (m, 1H), 2.52–2.41 (m, 1H), 2.41–2.33 (m, 1H) ppm (spectral data is consistent with reported values^32^).

**HRMS** (ESI^+^) calcd for C_16_H_20_N_6_O_5_ 376.1495 found 377.1567 (MH^+^)

### C13-O-3,4-(OH)_2_-Ph synthesis

3,4-Dihydroxybenzoate: Synthesized according to the above general procedure to yield the title product (30%).

**^1^H NMR** (600 MHz, D_2_O) δ 7.59 (d, J = 2.1 Hz, 1H), 7.55 (s, 1H), 7.02 (d, J = 8.4 Hz, 1H), 4.87 (s, 1H), 4.67–4.59 (m, 1H), 4.35–4.28 (m, 1H), 4.07–3.99 (m, 1H), 3.86–3.79 (m, 1H), 3.64–3.52 (m, 1H), 2.47–2.41 (m, 1H), 2.41–2.32 (m, 1H) ppm

**HRMS** (ESI^+^) calcd for C_16_H_20_N_6_O_6_ 392.1444 found 393.1513 (MH^+^)

### Thermofluor (TF) assay

Thermofluor assays for the STX congeners were performed as previously described^10,29^. Briefly, two-fold serial dilutions of each toxin were prepared in 150 mM NaCl, 10 mM HEPES, pH 7.4. 20 μL samples containing 1.1 μM *Rc*Sxph, *Rc*Sxph mutants, or *Np*Sxph, 5x SYPRO Orange dye (Sigma-Aldrich, cat. no. S5692, stock concentration 5,000x), 0-20 μM toxin, 150 mM NaCl, 10 mM HEPES, pH 7.4 were set up in 96-well PCR plates (Bio-Rad, cat. no. MLL9601), sealed with a microseal B adhesive sealing film (Bio-Rad, cat. no. MSB1001) and centrifuged (1 min, 230xg) prior to thermal denaturation using a CFX Connect Thermal Cycler (Bio-Rad, cat. no. 1855201). Fluorescence was measured using the HEX channel (excitation l=515-535 nm, emission l=560-580 nm). Samples were incubated at 25°C for 2 min followed by a temperature gradient from 25°C to 95°C at 0.2°C min^-1^, and final incubation at 95°C for 1 min.

For the toxin dose-response curves, Sxph melting temperatures (Tms) in the presence of varied toxin concentrations were calculated by fitting the denaturation curves using a Boltzmann function in GraphPad Prism (GraphPad Software) using the equation F=F_min_+(F_max_-F_min_)/(1+exp((Tm-T)/C)), where F is the fluorescence intensity at temperature T, F_min_ and F_max_ are the fluorescence intensities before and after the denaturation transition, respectively, Tm is the midpoint temperature of the transition, and C is the slope at Tm. ΔTms for Sxph in the absence (Tm_Sxph_) and presence (Tm_Sxph+Toxin_) of different toxin concentrations were calculated using the following equation: ΔTm=Tm_Sxph+Toxin-_Tm_Sxph_.

### Fluorescence polarization competition (FPc) assay

Fluorescence polarization competition (FPc) assays were performed as described^12^ using 100 µL total reaction volume per well and final concentrations of the fluorescent ligand and Sxph of: for *Rc*Sxph and *Rc*Sxph T563A, 1 nM F-STX and 12 nM *Rc*Sxph; for *Rc*Sxph Y558A, 1 nM F-STX and 2 nM *Rc*Sxph Y558A; for *Rc*Sxph F561A, 1 nM F-STX and 50 nM *Rc*Sxph F561A; for *Np*Sxph, 0.5 nM F-STX and 1.6 nM *Np*Sxph. For the displacement experiments, two-fold serial dilutions of unlabeled toxins were prepared using in a buffer of 150 mM NaCl, 10 mM HEPES, pH 7.4 at the concentration range of 0-1 µM. Samples containing F-STX, Sxph and 150 mM NaCl, 10 mM HEPES, pH 7.4 were incubated for 1 h at room temperature (23 ± 2°C) protected from light. Subsequently, 50 µL the equilibrated Sxph:F-STX solution was mixed with 50 µL of each toxin dilution series in 96-well black flat-bottomed polystyrene microplates (Greiner Bio-One, cat. no. 655900). The plate was sealed with an aluminum foil sealing film AlumaSeal II (Excel Scientific, cat. no. AF-100) and incubated at room temperature (23 ± 2°C) for 2 h to attain equilibrium. Measurements were performed at 25°C on a Synergy H1 microplate reader (BioTek) using the polarization filter setting (excitation l=485 nm, emission l=528 nm). Data were normalized using the following equation^29^: P = (P_Sxph:toxin_ − P_Toxin_)/(P_Ctrl_ − P_Toxin_), where P is the polarization measured at a given toxin concentration, P_Sxph:toxin_ is the polarization of Sxph:toxin mixture, P_Toxin_ is the polarization of toxin in the absence of Sxph, and P_Ctrl_ is the maximum polarization of Sxph in the absence of unlabeled toxin. The concentrations of the competing toxin causing displacement of 50% of bound F-STX (IC_50_) were calculated by fitting normalized fluorescence polarization as a function of toxin concentration using the nonlinear regression analysis in GraphPad Prism (GraphPad Software).

### Crystallization and structure determination

*Rc*Sxph and mutants thereof were crystallized as previously described for *Rc*Sxph with STX. Briefly, purified protein was exchanged into a buffer of 10 mM NaCl, 10 mM HEPES, pH 7.4 and concentrated to 65 mg mL^-1^ using a 50-kDa cutoff Amicon Ultra centrifugal filter unit (Millipore). Crystallization was set up by hanging drop vapor diffusion using a 24-well VDX plate with sealant (Hampton Research) using 3 μL drops having a 2:1 (v:v) ratio of protein and precipitant. For co-crystallization with STX or C13-OBz and the target *Rc*Sxph mutants were mixed in a molar ratio of 1:1.1 Sxph:toxin and incubated on ice for 1 hour before setting up crystallization. For toxin co-crystallization, *Rc*Sxph:C13-OBz, *Rc*Sxph Y558A:C13-OBz, *Rc*Sxph F561A:C13-OBz, *Rc*Sxph T563A:C13-OBz, and *Rc*Sxph F561A:STX were crystallized from solutions containing 20-33% (v/v) 2-methyl-2,4-pentanediol, 5 % (w/v) PEG 8000, 0.08-0.2 M sodium cacodylate, pH 6.5.

For *Np*Sxph crystallization, protein was SEC purified in 30 mM NaCl, 10 mM HEPES, pH 7.4 and concentrated to 30-40 mg mL^-1^ using a 50-kDa cutoff Amicon Ultra centrifugal filter unit (Millipore) as previously described. For co-crystallization with STX congeners, *Np*Sxph and C13-OBz or C13-NBz were mixed in a molar ratio of 1:1.2 *Np*Sxph:toxin and incubated on ice for 1 hour before setting up the crystallization trays. *Np*Sxph crystals were obtained by hanging drop vapor diffusion at 4°C using 1:1 (v/v) ratio of protein and precipitant from 400 nL drops set with Mosquito crystal (SPT Labtech) using 20-25% (v/v) PEG 400, 4-5% (w/v) PGA-LM, 100-200 mM sodium acetate, pH 5.0. *Np*Sxph: C13-OBz, and *Np*Sxph: C13-NBz crystals were harvested and flash-frozen in liquid nitrogen without additional cryoprotectant.

X-ray datasets for *Rc*Sxph:STX congener structures were collected at 100K at the Advanced Light Source (ALS) 8.3.1 beamline (Berkeley, CA). *Np*Sxph:STX congener complexes were collected at 100K at the Advanced Photon Source (APS) 23-ID-B beamline (Lemont, IL). Data were processed with XDS^42^ and scaled and merged with Aimless^43^. The Sxph:toxin structures were solved by molecular replacement using the apo-*Rc*Sxph structure (PDB:6O0D)^11^ or apo-*Np*Sxph structure (PDB: 8D6G)^10^ as a search model in Phaser from PHENIX^44^. The electron density map and the model were manually checked in COOT^45^ and iterative refinement was performed using phenix.refine ^44^. The quality of all models was assessed using MolProbity^46^ and refinement statistics.

## Acknowledgements

The authors thank K. Huang for technical help. The authors acknowledge the UCSF MSG X-ray Facility has supported by the UCSF Program for Breakthrough Biomedical Research and funded in part by the Sandler Foundation. Mass spectroscopic characterization of new STX derivatives utilized a Thermo Exploris 240 LC/MS system (RRID:SCR_022216) that was purchased with funding from Stanford C-ShaRP (RRID:SCR_022986) and was supported by Stanford University Mass Spectrometry (RRID:SCR_017801). This work was supported by DoD grants HDTRA-1-19-1-0040, HDTRA-1-21-1-0011, and HDTRA-1-23-1-0026 to D.L.M., and NIH-NIGMS R01-GM117263-05 to J.D.

## Author Contributions

S.Z., Z.C., J.D., and D.L.M. conceived the study and designed the experiments. S.Z. and Z.C. produced and purified Sxphs and performed the TF assay. S.Z. performed the FPc studies. S.Z. and Z.C. determined the crystal structures of Sxphs. E.R.P., R.G.B., and T.A.B. synthesized and characterized STX congeners. S.Z., Z.C., and D.L.M. analyzed data. J.D., and D.L.M. provided guidance and support. S.Z., Z.C., J.D., and D.L.M. wrote the paper.

## Data and materials availability

Coordinates and structure factors for *Rc*Sxph:C13-OBz (PDB:9YAR), *Rc*Sxph F561A (PDB:9YAV), *Rc*Sxph F561A:STX (PDB:9YBE), *Rc*Sxph F561A:C13-OBz (PDB:9YBD), *Rc*Sxph T563A (PDB:9YAT), *Rc*Sxph T563A:C13-OBz (PDB:9YBF), *Rc*Sxph Y558A:C13-OBz (PDB:9YAS), *Np*Sxph:C13-OBz (PDB:9Y92), and *Np*Sxph:C13-NBz (PDB:9Y91) are deposited with the RCSB and will be released upon publication. Requests for material should be sent to D.L.M.

## References

1. Thottumkara, A.P., Parsons, W.H., and Du Bois, J. (2014). Saxitoxin. Angew Chem Int Ed Engl 53, 5760–5784. 10.1002/anie.201308235.

2. Wiese, M., D’Agostino, P.M., Mihali, T.K., Moffitt, M.C., and Neilan, B.A. (2010). Neurotoxic alkaloids: saxitoxin and its analogs. Mar Drugs 8, 2185–2211. 10.3390/md8072185.

3. Smith, Z.J., Arlinghaus, K.M., Boyer, G.L., and Hapeman, C.J. (2025). A Fresh Perspective on Cyanobacterial Paralytic Shellfish Poisoning Toxins: History, Methodology, and Toxicology. Mar Drugs 23. 10.3390/md23070271.

4. Deng, H., Shang, X., Zhu, H., Huang, N., Wang, L., and Sun, M. (2025). Saxitoxin: A Comprehensive Review of Its History, Structure, Toxicology, Biosynthesis, Detection, and Preventive Implications. Mar Drugs 23. 10.3390/md23070277.

5. Gribble, M.O., Bennett, B.J., Liddie, J.M., Borchert, W., Pfluger, B.A., Segars, J.S., Keast, J.M., Hans, A., Kikkeri, N.S., Shin, C., et al. (2025). Global epidemiology of paralytic shellfish poisoning: a systematic search literature review. Lancet Planet Health 9, 101271. 10.1016/j.lanplh.2025.05.001.

6. Duran-Riveroll, L.M., and Cembella, A.D. (2017). Guanidinium Toxins and Their Interactions with Voltage-Gated Sodium Ion Channels. Mar Drugs 15. 10.3390/md15100303.

7. Llewellyn, L.E. (2006). Saxitoxin, a toxic marine natural product that targets a multitude of receptors. Nat Prod Rep 23, 200–222. 10.1039/b501296c.

8. Shen, H., Liu, D., Wu, K., Lei, J., and Yan, N. (2019). Structures of human Nav1.7 channel in complex with auxiliary subunits and animal toxins. Science 363, 1303–1308. 10.1126/science.aaw2493.

9. Shen, H., Li, Z., Jiang, Y., Pan, X., Wu, J., Cristofori-Armstrong, B., Smith, J.J., Chin, Y.K.Y., Lei, J., Zhou, Q., et al. (2018). Structural basis for the modulation of voltage-gated sodium channels by animal toxins. Science 362. 10.1126/science.aau2596.

10. Chen, Z., Zakrzewska, S., Hajare, H.S., Alvarez-Buylla, A., Abderemane-Ali, F., Bogan, M., Ramirez, D., O’Connell, L.A., Du Bois, J., and Minor, D.L., Jr. (2022). Definition of a saxitoxin (STX) binding code enables discovery and characterization of the anuran saxiphilin family. Proc Natl Acad Sci U S A 119, e2210114119. 10.1073/pnas.2210114119.

11. Yen, T.-J., Lolicato, M., Thomas-Tran, R., Du Bois, J., and Minor, D.L., Jr., (2019). Structure of the Saxiphilin:saxitoxin (STX) complex reveals a convergent molecular recognition strategy for paralytic toxins. Sci Adv 5. 10.1126/sciadv.aax2650.

12. Zakrzewska, S., Nixon, S.A., Chen, Z., Hajare, H.S., Park, E.R., Mulcahy, J.V., Arlinghaus, K.M., Neu, E., Konovalov, K., Provasi, D., et al. (2025). Structural basis for saxitoxin congener binding and neutralization by anuran saxiphilins. Nat Commun 16, 3885. 10.1038/s41467-025-58903-2.

13. Davio, S.R. (1985). Neutralization of saxitoxin by anti-saxitoxin rabbit serum. Toxicon 23, 669–675. 10.1016/0041-0101(85)90371-x.

14. Benton, B.J., Rivera, V.R., Hewetson, J.F., and Chang, F.C. (1994). Reversal of saxitoxin-induced cardiorespiratory failure by a burro-raised alpha-STX antibody and oxygen therapy. Toxicol Appl Pharmacol 124, 39–51. 10.1006/taap.1994.1006.

15. Kaufman, B., Wright, D.C., Ballou, W.R., and Monheit, D. (1991). Protection against tetrodotoxin and saxitoxin intoxication by a cross-protective rabbit anti-tetrodotoxin antiserum. Toxicon 29, 581–587. 10.1016/0041-0101(91)90052-s.

16. Wang, S., Wang, D., Shen, W.T., Kai, M., Yu, Y., Peng, Y., Xian, N., Fang, R.H., Gao, W., and Zhang, L. (2024). Protein-Loaded Cellular Nanosponges for Dual-Biomimicry Neurotoxin Countermeasure. Small 20, e2309635. 10.1002/smll.202309635.

17. Nixon, S.A., Zakrzewska, S., Jang, S., Huang, K., Bara, A., Chen, Z., Goss, D.R., Park, E.R., Du Bois, J., and Minor, D.L. Jr., (2025). Saxiphilin functions as a ‘toxin sponge’ protein that counteracts the effects of saxitoxin poisoning. bioRxiv. 10.1101/2025.11.20.689596.

18. Ondrus, A.E., Lee, H.L., Iwanaga, S., Parsons, W.H., Andresen, B.M., Moerner, W.E., and Du Bois, J. (2012). Fluorescent saxitoxins for live cell imaging of single voltage-gated sodium ion channels beyond the optical diffraction limit. Chem Biol 19, 902–912. 10.1016/j.chembiol.2012.05.021.

19. Thomas-Tran, R., and Du Bois, J. (2016). Mutant cycle analysis with modified saxitoxins reveals specific interactions critical to attaining high-affinity inhibition of hNaV1.7. Proc Natl Acad Sci U S A 113, 5856–5861. 10.1073/pnas.1603486113.

20. Elleman, A.V., Milicic, N., Williams, D.J., Simko, J., Liu, C.J., Haynes, A.L., Ehrlich, D.E., Makinson, C.D., and Du Bois, J. (2024). Behavioral control through the direct, focal silencing of neuronal activity. Cell Chem Biol 31, 1324–1335 e1320. 10.1016/j.chembiol.2024.04.003.

21. Park, E.R., Denomme, N., Hajare, H.S., and Du Bois, J. (2025). A chemogenetic ligand-receptor pair for voltage-gated sodium channel subtype-selective inhibition. bioRxiv. 10.1101/2025.11.03.686046.

22. Guo, Y., Li, Y., Chen, S., Wu, Y., Poll, O., Ren, Z., Liu, Z., Vlkolinsky, R., Bajo, M., Prier, C.K., et al. (2025). Scalable total synthesis of saxitoxin and related natural products. Nature 646, 351–357. 10.1038/s41586-025-09551-5.

23. Hoehne, A., Behera, D., Parsons, W.H., James, M.L., Shen, B., Borgohain, P., Bodapati, D., Prabhakar, A., Gambhir, S.S., Yeomans, D.C., et al. (2013). A 18F-labeled saxitoxin derivative for in vivo PET-MR imaging of voltage-gated sodium channel expression following nerve injury. J Am Chem Soc 135, 18012–18015. 10.1021/ja408300e.

24. Pajouhesh, H., Delwig, A., Beckley, J.T., Klas, S., Monteleone, D., Zhou, X., Luu, G., Du Bois, J., Hunter, J.C., and Mulcahy, J.V. (2022). Discovery of Selective Inhibitors of Na(V)1.7 Templated on Saxitoxin as Therapeutics for Pain. ACS Med Chem Lett 13, 1763–1768. 10.1021/acsmedchemlett.2c00378.

25. Pajouhesh, H., Beckley, J.T., Delwig, A., Hajare, H.S., Luu, G., Monteleone, D., Zhou, X., Ligutti, J., Amagasu, S., Moyer, B.D., et al. (2020). Discovery of a selective, state-independent inhibitor of Na(V)1.7 by modification of guanidinium toxins. Sci Rep 10, 14791. 10.1038/s41598-020-71135-2.

26. Mahar, J., Lukacs, G.L., Li, Y., Hall, S., and Moczydlowski, E. (1991). Pharmacological and biochemical properties of saxiphilin, a soluble saxitoxin-binding protein from the bullfrog (Rana catesbeiana). Toxicon 29, 53–71.

27. Llewellyn, L.E., and Moczydlowski, E.G. (1994). Characterization of saxitoxin binding to saxiphilin, a relative of the transferrin family that displays pH-dependent ligand binding. Biochemistry 33, 12312–12322.

28. Abderemane-Ali, F., Rossen, N.D., Kobiela, M.E., Craig, R.A., Garrison, C.E., Chen, Z., Colleran, C.M., O’Connell, L.A., Du Bois, J., Dumbacher, J.P., and Minor, D.L. (2021). Evidence that toxin resistance in poison birds and frogs is not rooted in sodium channel mutations and may rely on “toxin sponge” proteins. J Gen Physiol 153. 10.1085/jgp.202112872.

29. Chen, Z., Zakrzewska, S., Hajare, H.S., Du Bois, J., and Minor, D.L.Jr., (2023). Expression, purification, and characterization of anuran saxiphilins using thermofluor, fluorescence polarization, and isothermal titration calorimetry. STAR Protoc 5, 102792. 10.1016/j.xpro.2023.102792.

30. Akimoto, T., Masuda, A., Yotsu-Yamashita, M., Hirokawa, T., and Nagasawa, K. (2013). Synthesis of saxitoxin derivatives bearing guanidine and urea groups at C13 and evaluation of their inhibitory activity on voltage-gated sodium channels. Org Biomol Chem 11, 6642–6649. 10.1039/c3ob41398e.

31. Onodera, H., Satake, M., Oshima, Y., Yasumoto, T., and Carmichael, W.W. (1997). New saxitoxin analogues from the freshwater filamentous cyanobacterium Lyngbya wollei. Nat Toxins 5, 146–151. 10.1002/1522-7189(1997)5:4<146::AID-NT4>3.0.CO;2-V.

32. Negri, A., Stirling, D., Quilliam, M., Blackburn, S., Bolch, C., Burton, I., Eaglesham, G., Thomas, K., Walter, J., and Willis, R. (2003). Three novel hydroxybenzoate saxitoxin analogues isolated from the dinoflagellate Gymnodinium catenatum. Chem Res Toxicol 16, 1029–1033. 10.1021/tx034037j.

33. Vale, P. (2008). Complex profiles of hydrophobic paralytic shellfish poisoning compounds in Gymnodinium catenatum identified by liquid chromatography with fluorescence detection and mass spectrometry. J Chromatogr A 1195, 85–93. 10.1016/j.chroma.2008.04.073.

34. Niesen, F.H., Berglund, H., and Vedadi, M. (2007). The use of differential scanning fluorimetry to detect ligand interactions that promote protein stability. Nat Protoc 2, 2212–2221. 10.1038/nprot.2007.321.

35. Huynh, K., and Partch, C.L. (2015). Analysis of protein stability and ligand interactions by thermal shift assay. Curr Protoc Protein Sci 79, 28 29 21-14. 10.1002/0471140864.ps2809s79.

36. Rossi, A.M., and Taylor, C.W. (2011). Analysis of protein-ligand interactions by fluorescence polarization. Nat Protoc 6, 365–387. 10.1038/nprot.2011.305.

37. Lee, H., Lolicato, M., Arrigoni, C., and Minor, D.L., Jr. (2021). Production of K(2P)2.1 (TREK-1) for structural studies. Methods Enzymol 653, 151–188. 10.1016/bs.mie.2021.02.013.

38. Andresen, B.M., and Du Bois, J. (2009). De novo synthesis of modified saxitoxins for sodium ion channel study. J Am Chem Soc 131, 12524–12525. 10.1021/ja904179f.

39. Mulcahy, J.V., and Du Bois, J. (2008). A stereoselective synthesis of (+)-gonyautoxin 3. Journal of the American Chemical Society 130, 12630-+. 10.1021/ja805651g.

40. Beutner, G.L., Young, I.S., Davies, M.L., Hickey, M.R., Park, H., Stevens, J.M., and Ye, Q. (2018). TCFH-NMI: Direct Access to N-Acyl Imidazoliums for Challenging Amide Bond Formations. Org Lett 20, 4218–4222. 10.1021/acs.orglett.8b01591.

41. Walker, J.R., Merit, J.E., Thomas-Tran, R., Tang, D.T.Y., and Du Bois, J. (2019). Divergent Synthesis of Natural Derivatives of (+)-Saxitoxin Including 11-Saxitoxinethanoic Acid. Angew Chem Int Ed Engl 58, 1689–1693. 10.1002/anie.201811717.

42. Kabsch, W. (2010). Xds. Acta Crystallogr D Biol Crystallogr 66, 125–132. 10.1107/S0907444909047337.

43. Evans, P.R., and Murshudov, G.N. (2013). How good are my data and what is the resolution? Acta Crystallogr D Biol Crystallogr 69, 1204–1214. 10.1107/S0907444913000061.

44. Adams, P.D., Afonine, P.V., Bunkoczi, G., Chen, V.B., Davis, I.W., Echols, N., Headd, J.J., Hung, L.W., Kapral, G.J., Grosse-Kunstleve, R.W., et al. (2010). PHENIX: a comprehensive Python-based system for macromolecular structure solution. Acta Crystallogr D Biol Crystallogr 66, 213–221. 10.1107/S0907444909052925.

45. Emsley, P., and Cowtan, K. (2004). Coot: model-building tools for molecular graphics. Acta Crystallogr D Biol Crystallogr 60, 2126–2132.

46. Williams, C.J., Headd, J.J., Moriarty, N.W., Prisant, M.G., Videau, L.L., Deis, L.N., Verma, V., Keedy, D.A., Hintze, B.J., Chen, V.B., et al. (2018). MolProbity: More and better reference data for improved all-atom structure validation. Protein Sci 27, 293–315. 10.1002/pro.3330.

