## Supplementary Figures S1-S4 and Table S1 for "Saxiphilin is a broad-spectrum toxin sponge for C13-modified saxitoxins"

10-Mar-26

[7]Molecular Biophysics and Integrated Bio-imaging Division
Lawrence Berkeley National Laboratory, Berkeley, CA 94720 USA

[2]Department of Chemistry
Stanford University, Stanford, CA 94305 USA

[†] Present address:
Department of Anatomy and Physiology
Shanghai Jiao Tong University School of Medicine
Shanghai, 200025, China

### Figure S1

Zakrzewska *et al.*

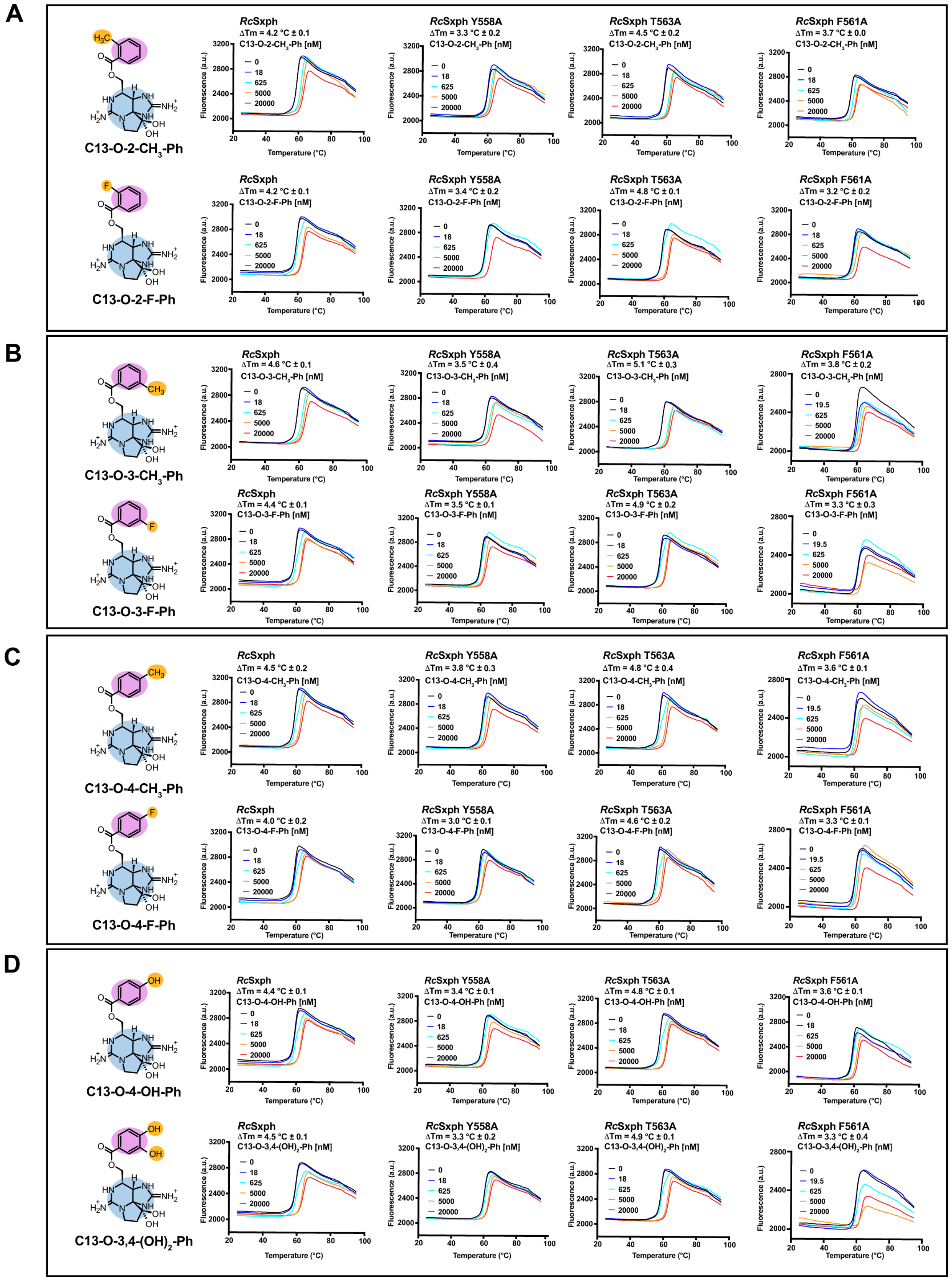

**Figure S1 TF assays show binding preferences for *Rc*Sxph and mutants.** Exemplar TF assay results for *Rc*Sxph, *R*cSxph Y558A, *R*cSxph T563A, and *R*cSxph F561A with **A,** C13-O-2-$CH_3$-Ph and C13-O-2-F-Ph, **B,** C13-O-3-$CH_3$-Ph and C13-O-3-F-Ph, **C,** C13-O-4-$CH_3$-Ph and C13-O-4-F-Ph, and **D,** C13-O-4-OH-Ph and C13-O-3,4-$(OH)_2$-Ph at 0 nM (black), 18 nM (orange), 625 nM (green), 5000 nM (magenta), and 20000 nM (red).

**Figure S2** **Zakrzewska *et al.***

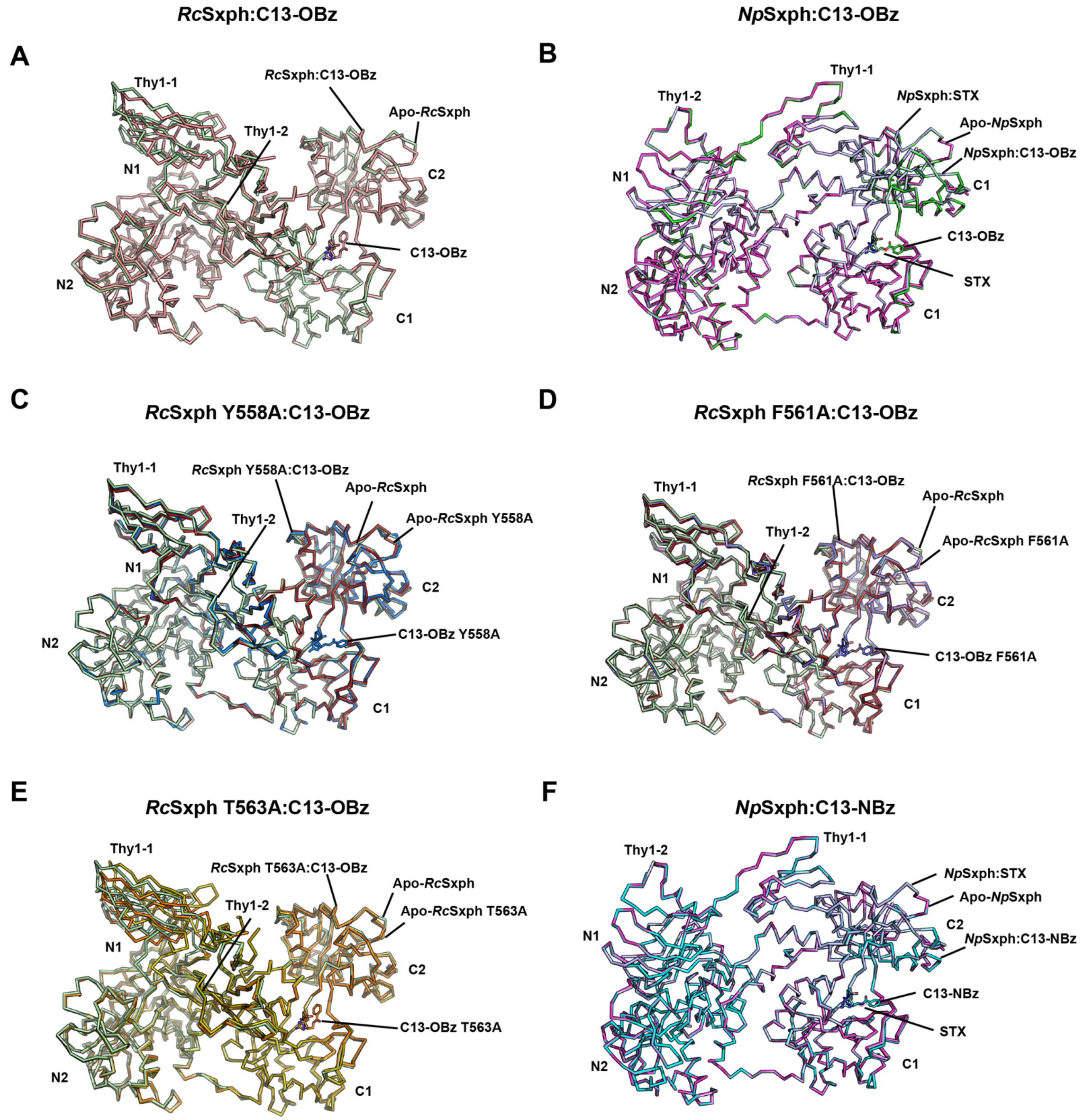


**Figure S2 Structural comparisons of C13-STX congener complexes with apo- and STX-bound Sxphs.** **A,** *Rc*Sxph:C13-OBz (salmon) and apo-*Rc*Sxph (6O0D) (pale green)(*1*). **B,** *Np*Sxph:C13-OBz (green), apo-

*Np*Sxph (magenta) (8D6G)(*2*) and *Np*Sxph:STX (light blue) (8D6M)(*2*). **C,** *Rc*Sxph Y558A:C13-OBz (marine), apo-*Rc*Sxph Y558A (8D6P) (firebrick)(*2*), and apo-*Rc*Sxph (6O0D) (pale green)(*1*). **D,** *Rc*Sxph F561A:C13-OBz (slate), apo-*Rc*Sxph F561A (pink), and apo-*Rc*Sxph (6O0D) (pale green)(*1*). **E,** *Rc*Sxph T563A:C13-OBz (orange), apo-*Rc*Sxph T563A (olive), and apo-*Rc*Sxph (6O0D) (pale green)(*1*). **F,** *Np*Sxph:C13-NBz (cyan), apo-*Np*Sxph (magenta) (8D6G)(*2*) and *Np*Sxph:STX (light blue) (8D6M)(*2*). C13-OBz, C13-NBz, and STX are shown as sticks.

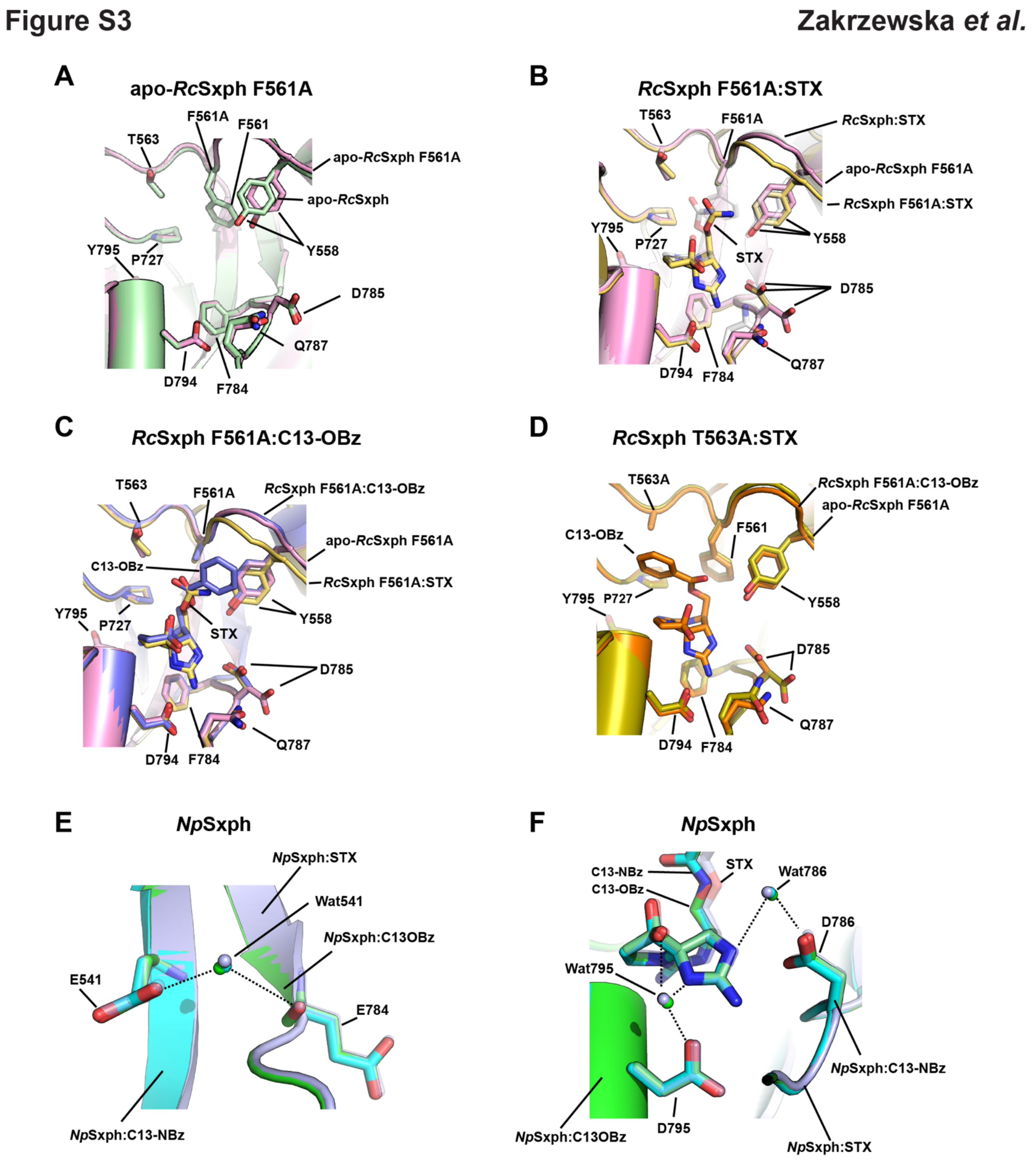


**Figure S3 Sxph binding pocket comparisons and water networks.** Binding pocket comparisons for **A,** apo-*Rc*Sxph F561A (pink) and apo-*Rc*Sxph (6O0D) (pale green)(*1*). **B,** apo-*Rc*Sxph F561A (pink), *Rc*Sxph

F561A:STX (yellow), and *Rc*Sxph:STX (6O0F) (white)(*1*). **C,** apo-*Rc*Sxph F561A (pink), *Rc*Sxph F561A:STX (yellow), and *Rc*Sxph F561A:C13-OBz (slate). And **D,** *Rc*Sxph T563A:C13-OBz (orange) and apo-*Rc*Sxph T563A (olive). **E,** and **F,** Water interactions for **E,** Wat541 and **F,** Wat786 and Wat795 in *Np*Sxph:C13-OBz (green), *Np*Sxph:C13-NBz (cyan), and *Np*Sxph:STX (light blue) (8D6M)(*2*).

**Figure S4** **Zakrzewska *et al.***

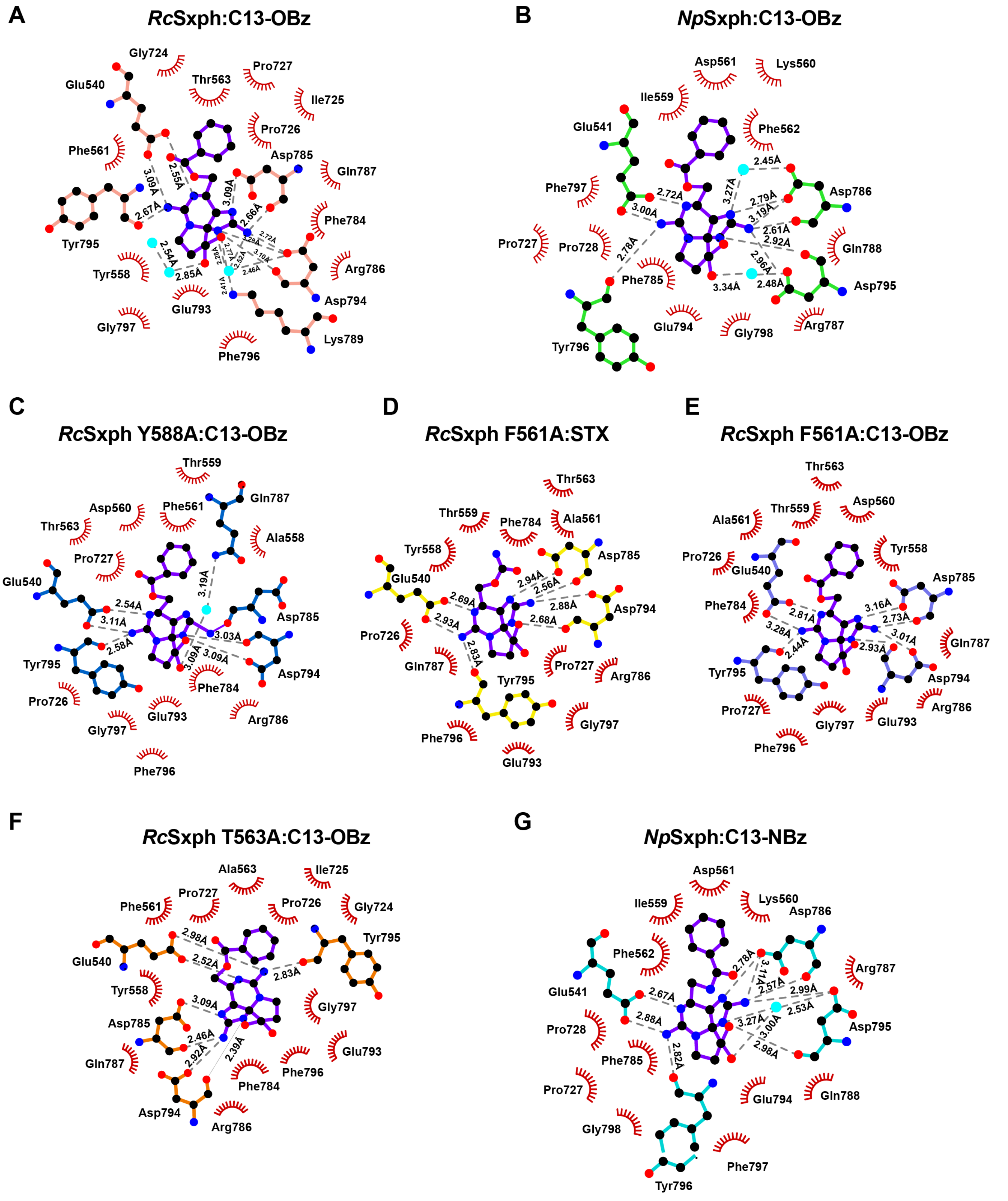

**Figure S4 Sxph:STX congener interactions A-G,** LIGPLOT(*3*) diagrams showing interactions (5.0Å cutoff) for **A,** *Rc*Sxph:C13-OBz (salmon), **B,** *Np*Sxph:C13-OBz (green), **C,** *Rc*Sxph Y558A:C13-OBz (marine), **D,** *Rc*Sxph F561A:STX (yellow). **E,** *Rc*Sxph F561A:C13-OBz (slate), **F,** *Rc*Sxph T563A:C13-OBz (orange), **G,** *Np*Sxph:C13-NBz (cyan). Toxin is purple in all panels.

**Table S1 Crystallographic data collection and refinement statistics**

| | ***Rc*Sxph:C13-OBz (PDB:9YAR)** | ***Rc*Sxph F561A (PDB:9YAV)** | ***Rc*Sxph F561A:STX (PDB:9YBE)** | ***Rc*Sxph F561A:C13-OBz (PDB:9YBD)** | ***Rc*Sxph T563A (PDB:9YAT)** |
|---|---|---|---|---|---|
| **Data Collection** | | | | | |
| Space group | $P2_12_12_1$ | $P2_12_12_1$ | $P2_12_12_1$ | $P2_12_12_1$ | $P2_12_12_1$ |
| Cell dimensions a/b/c (Å) | 95.98, 105.69, 253.28 | 96.35, 110.24, 254.70 | 96.06, 105.11, 253.26 | 96.42, 110.04, 256.01 | 96.30, 109.69, 253.14 |
| α/β/γ (°) | 90, 90, 90 | 90, 90, 90 | 90, 90, 90 | 90, 90, 90 | 90, 90, 90 |
| Resolution (Å) | 48.77-2.40 (2.43-2.40) | 48.17-2.23 (2.26-2.23) | 48.54-2.35 (2.38-2.35) | 48.21-2.40 (2.43-2.40) | 47.66-2.50 (2.53-2.50) |
| Rmerge (%) | 0.0246 (0.9399) | 0.01606 (0.5350) | 0.01907 (0.6267) | 0.02175 (0.7213) | 0.02094 (0.6056) |
| I / σ I | 15.9(0.8) | 19.6 (1.3) | 17.5 (1.3) | 17.3 (1.1) | 14.6 (1.3) |
| CC(1/2) | 1.000 (0.391) | 1.000 (0.674) | 0.999 (0.599) | 1.000 (0.668) | 0.999 (0.618) |
| Completeness (%) | 100.00 (100.00) | 100.00 (100.00) | 100.00 (100.00) | 100.00 (99.90) | 99.2 (97.9) |
| Redundancy | 13.3 (13.8) | 13.1 (13.3) | 13.2 (13.0) | 13.3 (14.0) | 13.0 (14.1) |
| Total reflections | 1347412 (67836) | 1745597 (86542) | 1423562 (68105) | 1425339 (72920) | 1201741 (63639) |
| Unique reflections | 101479 (4928) | 132748 (6507) | 107521 (5258) | 107174 (5220) | 92708 (4526) |
| Wilson B-factor | 64.45 | 55.66 | 59.98 | 61.69 | 65.33 |
| Wavelength (Å) | 1.116 | 1.116 | 1.116 | 1.116 | 1.116 |
| **Refinement** | | | | | |
| $R_{work}$ / $R_{free}$ (%) | 24.95/27.82 | 21.03/24.38 | 23.96/27.44 | 22.93/25.86 | 20.68/24.58 |
| No. of chains in AU | 2 | 2 | 2 | 2 | 2 |
| No. of protein atoms | 12630 | 12737 | 12733 | 12748 | 12751 |
| No. of ligand atoms | 52 | 0 | 42 | 52 | 0 |
| No. of water atoms | 329 | 370 | 286 | 62 | 171 |
| RMSD bond lengths (Å) | 0.004 | 0.005 | 0.003 | 0.005 | 0.003 |
| RMSD angles (°) | 0.62 | 0.68 | 0.57 | 0.71 | 0.55 |
| Ramachandran favored/allowed/outliers (%) | 94.66/4.91/0.43 | 95.92/3.83/0.24 | 95.26/4.37/0.36 | 94.90/4.79/0.30 | 95.93/3.83/0.24 |

**Table S1 Crystallographic data collection and refinement statistics**

| | ***Rc*Sxph T563A:C13-OBz (PDB:9YBF)** | ***Rc*Sxph Y558A:C13-OBz (PDB:9YAS)** | ***Np*Sxph:C13-OBz (PDB:9Y92)** | ***Np*Sxph:C13-NBz (PDB:9Y91)** |
|---|---|---|---|---|
| **Data Collection** | | | | |
| Space group | $P2_12_12_1$ | $P2_12_12_1$ | R3 | R3 |
| Cell dimensions a/b/c (Å) | 96.12, 105.71, 251.74 | 96.20, 109.33, 254.69 | 228.64, 228.64, 67.18 | 228.98, 228.98, 67.48 |
| α/β/γ (°) | 90, 90, 90 | 90, 90, 90 | 90, 90, 120 | 90, 90, 120 |
| Resolution (Å) | 48.73-2.45 (2.48-2.45) | 48.10-2.45 (2.48-2.45) | 43.21-1.90 (1.92-1.90) | 43.27-1.95 (1.97-1.95) |
| Rmerge (%) | 0.02232 (0.6200) | 0.02061 (0.6854) | 0.05688 (1.858) | 0.08223 (2.659) |
| I / σ I | 14.9 (1.2) | 18.3 (1.1) | 16.01 (1.15) | 12.65 (0.81) |
| CC(1/2) | 1.000 (0.526) | 1.000 (0.636) | 0.999 (0.562) | 0.999 (0.286) |
| Completeness (%) | 99.3 (99.0) | 100.00 (100.00) | 99.93 (100.00) | 99.93 (99.63) |
| Redundancy | 13.2 (14.0) | 13.3 (13.7) | 10.4 (10.8) | 10.7 (9.8) |
| Total reflections | 1246248 (64912) | 1319377 (66866) | 1073550 (37791) | 1024673 (31869) |
| Unique reflections | 94340 (4624) | 99513 (4889) | 103144 (3488) | 96135 (3245) |
| Wilson B-factor | 61.55 | 63.43 | 46.73 | 46.31 |
| Wavelength (Å) | 1.116 | 1.116 | 1.033 | 1.033 |
| **Refinement** | | | | |
| $R_{work}$ / $R_{free}$ (%) | 23.15/26.26 | 24.94/28.39 | 19.39/22.33 | 19.41/22.28 |
| No. of chains in AU | 2 | 2 | 1 | 1 |
| No. of protein atoms | 12688 | 12616 | 6377 | 6384 |
| No. of ligand atoms | 52 | 52 | 42 | 42 |
| No. of water atoms | 200 | 213 | 400 | 400 |
| RMSD bond lengths (Å) | 0.004 | 0.005 | 0.011 | 0.009 |
| RMSD angles (°) | 0.69 | 0.69 | 0.95 | 0.93 |
| Ramachandran favored/allowed/outliers (%) | 95.11/4.46/0.43 | 94.16/5.10/0.74 | 96.06/3.81/0.12 | 95.57/4.18/0.25 |

#### References

1. T.-J. Yen, M. Lolicato, R. Thomas-Tran, J. Du Bois, D. L. Minor, Jr., Structure of the Saxiphilin:saxitoxin (STX) complex reveals a convergent molecular recognition strategy for paralytic toxins. *Sci Adv* **5**, (2019).

2. Z. Chen *et al.*, Definition of a saxitoxin (STX) binding code enables discovery and characterization of the anuran saxiphilin family. *Proc Natl Acad Sci U S A* **119**, e2210114119 (2022).

3. A. C. Wallace, R. A. Laskowski, J. M. Thornton, LIGPLOT: a program to generate schematic diagrams of protein-ligand interactions. *Protein Eng* **8**, 127-134 (1995).
